# Transcriptomic analyses implicate neuronal plasticity and chloride homeostasis in ivermectin resistance and recovery in a parasitic nematode

**DOI:** 10.1101/2021.11.12.468372

**Authors:** Roz Laing, Stephen R. Doyle, Jennifer McIntyre, Kirsty Maitland, Alison Morrison, David J. Bartley, Ray Kaplan, Umer Chaudhry, Neil Sargison, Andy Tait, James A. Cotton, Collette Britton, Eileen Devaney

## Abstract

The antiparasitic drug ivermectin plays an essential role in human and animal health globally. However, ivermectin resistance is widespread in veterinary helminths and there are growing concerns of sub-optimal responses to treatment in related helminths of humans. Despite decades of research, the genetic mechanisms underlying ivermectin resistance are poorly understood in parasitic helminths. This reflects significant uncertainty regarding the mode of action of ivermectin in parasitic helminths, and the genetic complexity of these organisms; parasitic helminths have large, rapidly evolving genomes and differences in evolutionary history and genetic background can confound comparisons between resistant and susceptible populations. We undertook a controlled genetic cross of a multi-drug resistant and a susceptible reference isolate of *Haemonchus contortus*, an economically important gastrointestinal nematode of sheep, and ivermectin-selected the F2 population for comparison with an untreated F2 control. RNA-seq analyses of male and female adults of all populations identified high transcriptomic differentiation between parental isolates, which was significantly reduced in the F2, allowing differences associated specifically with ivermectin resistance to be identified. In all resistant populations, there was constitutive upregulation of a single gene, *HCON_00155390:cky-1*, a putative pharyngeal-expressed transcription factor, in a narrow locus on chromosome V previously shown to be under ivermectin selection. In addition, we detected sex-specific differences in gene expression between resistant and susceptible populations, including constitutive upregulation of a P-glycoprotein, *HCON_00162780:pgp-11*, in resistant males only. After ivermectin selection, we identified differential expression of genes with roles in neuronal function and chloride homeostasis, which is consistent with an adaptive response to ivermectin-induced hyperpolarisation of neuromuscular cells. Overall, we show the utility of a genetic cross to identify differences in gene expression that are specific to ivermectin selection and provide a framework to better understand ivermectin resistance and recovery in parasitic helminths.

**Author Summary:** Parasitic helminths (worms) infect people and animals throughout the world and are largely controlled with mass administration of anthelmintic drugs. There are a very limited number of anthelmintics available and parasitic helminths can rapidly develop resistance to these drugs. Ivermectin is a widely used anthelmintic in both humans and animals, but resistance is now widespread in the veterinary field. We crossed ivermectin resistant and ivermectin susceptible parasitic helminths and treated them with ivermectin or left them as untreated controls. This provided resistant and susceptible populations with a similar genetic background with which to study differences in gene expression associated with ivermectin resistance. We identified upregulation of a gene with no previous association with drug resistance (*HCON_00155390:cky-1*) in male and female worms in all resistant populations. This gene is thought to be expressed in the helminth pharynx (mouthpart) and, in mammals, plays a role in controlling nerve function and protecting nerves from damage. This is consistent with the known effects of ivermectin in inhibiting helminth feeding through pharyngeal paralysis and implicates a novel mechanism that allows resistant worms to survive treatment.

## Introduction

Ivermectin, a ‘wonder drug’ which won its discoverers the Nobel Prize in Physiology or Medicine, is used primarily to control parasitic disease in humans and animals. Hundreds of millions of people are treated with ivermectin every year to control the filarial nematodes responsible for onchocerciasis (river blindness) and, in combination with albendazole, lymphatic filariasis [1]. Concurrently, ivermectin is one of the top selling animal health products in the world, used extensively in livestock to treat all major gastrointestinal nematodes, in addition to various ectoparasites, and for heartworm prophylaxis in domestic pets. However, ivermectin resistance is now widespread in the veterinary field [2, 3] and there are growing concerns of sub-optimal efficacy in humans [4-6].

It is hard to overstate the importance of ivermectin in human and animal health, yet there is limited understanding of either the molecular targets of ivermectin in parasitic worms or the mechanisms underlying ivermectin resistance (these may or may not be related). While candidate gene studies have identified single nucleotide polymorphisms (SNPs) or differences in expression of individual genes in resistant and susceptible parasites [7-11], these experiments do not account for differences in the evolutionary histories or genetic backgrounds of the resistant and susceptible populations, so the relevance of these genes to ivermectin resistance remains unclear. Further, exposure to an anthelmintic is likely to have wide-ranging biological effects in parasitic worms, yet the broad scale transcriptional response to drug treatment, and how this translates to the evolution of resistance, is largely unknown.

*Haemonchus contortus* is a highly pathogenic and economically important gastrointestinal nematode of small ruminants, and ivermectin resistant populations are now common throughout the world [3]. Relative to other parasitic nematodes, *H. contortus* is a tractable model for anthelmintic resistance research: it resides in the same phylogenetic group as the free-living model nematode *Caenorhabditis elegans* [12, 13] and has a reference quality genome assembly and annotation [14] allowing robust genome-wide analyses [15, 16].

To facilitate interrogation of complex traits, such as drug resistance, while controlling for between-population diversity, a genetic cross between genetically divergent populations differing in traits of interest can be generated [15, 17]. Progeny of the cross provide a controlled (admixed) population in which to identify genetic variants that co-vary with the trait of interest, for example after drug selection. Applying these principles, we performed a genetic cross between the *H. contortus* reference genome isolate (MHco3(ISE)) and a multi-drug resistant isolate (MHco18(UGA2004)) and selected adult worms of the F2 generation with ivermectin, benzimidazole or levamisole *in vivo* [18]. Genomic analyses of the F3 progeny pre- and post-treatment identified discrete loci under selection by each anthelmintic [18] and significantly narrowed a previously identified major quantitative trait locus (QTL) for ivermectin resistance on chromosome V [17].

In this paper, we characterise the broad scale transcriptomic response to ivermectin selection in populations with the same genetic background: male and female adults of the F2 generation of the genetic cross with and without ivermectin treatment. By also measuring differential expression in the two parental isolates, we identify genes specifically associated with ivermectin resistance (i.e. differentially expressed in both the ivermectin-treated genetic cross and the untreated resistant parent) and genes involved in the response to ivermectin treatment (i.e. differentially expressed in the ivermectin-treated genetic cross but not the untreated resistant parent). We highlight a single gene, *HCON_00155390*:*cky-1*, in the major QTL for ivermectin resistance, that is constitutively upregulated in all resistant populations, and a P-glycoprotein, *HCON_00162780:pgp-11*, that is upregulated in resistant males only. Several genes involved in neuronal development and regeneration, neuropeptide signalling and chloride homeostasis were also differentially expressed, providing a framework to better understand the effects of ivermectin on parasitic nematodes.

## Results

### High transcriptomic differentiation between parental isolates with admixture in the genetic cross

We generated RNA-seq data from male and female worms from both parental isolates (MHco3(ISE); drug susceptible reference isolate, referred to from here on as ‘MHco3’ and MHco18(UGA2004); multi-drug resistant and referred to from here on as ‘MHco18’) and from the F2 generation of the genetic cross after ivermectin treatment (F2IVM) and without treatment (F2CTL) (Figure S1). We also collected samples from the F2 after treatment with benzimidazole (F2BZ) for comparison. An average of 72 million (SD = 27.5 million) 150 bp paired-end reads were generated per sample (Figure S2A). Between 75 and 82% of MHco18 and MHco3 reads mapped to the reference (MHco3) genome, respectively, with the F2 samples showing intermediate mapping percentages (Figure S2B). This is consistent with the expected levels of polymorphism in each population relative to MHco3, with lower mapping percentages as populations diverge from the reference isolate [18]. Transcriptomic differentiation between isolates, treatment groups, and replicates was assessed using principal component analysis (PCA) and Poisson distance (Figure 1). Samples clustered well by isolate or treatment group for all male samples (Figures 1A and 1C); as expected, the parental isolates were highly differentiated (1A, PC1: 38% variance) and the genetic cross F2 samples were an admixture of the two. For the female samples (Figures 1B and 1D), the parental isolates were again well differentiated, but there was greater variance among all samples (1B, PC1: 53%) with less defined clustering of replicates and treatment groups. Differences between replicates broadly corresponded to the sample processing batch (highlighted by shape of point in 1B and sample name in 1D); this batch effect was included as a secondary factor in the DESeq2 analysis. The female F2CTL group was an outlier in the dataset (Figure S3), so was excluded from analysis. The specific cause of this variance is unknown: the cluster of genes (n = 129) that differentiated female F2CTL samples and the first replicate of all female samples were mostly uncharacterised; 16 had *C. elegans* homologues and in this subset there was no enrichment for expression in a particular tissue, or for any GO term.

**Figure 1.**
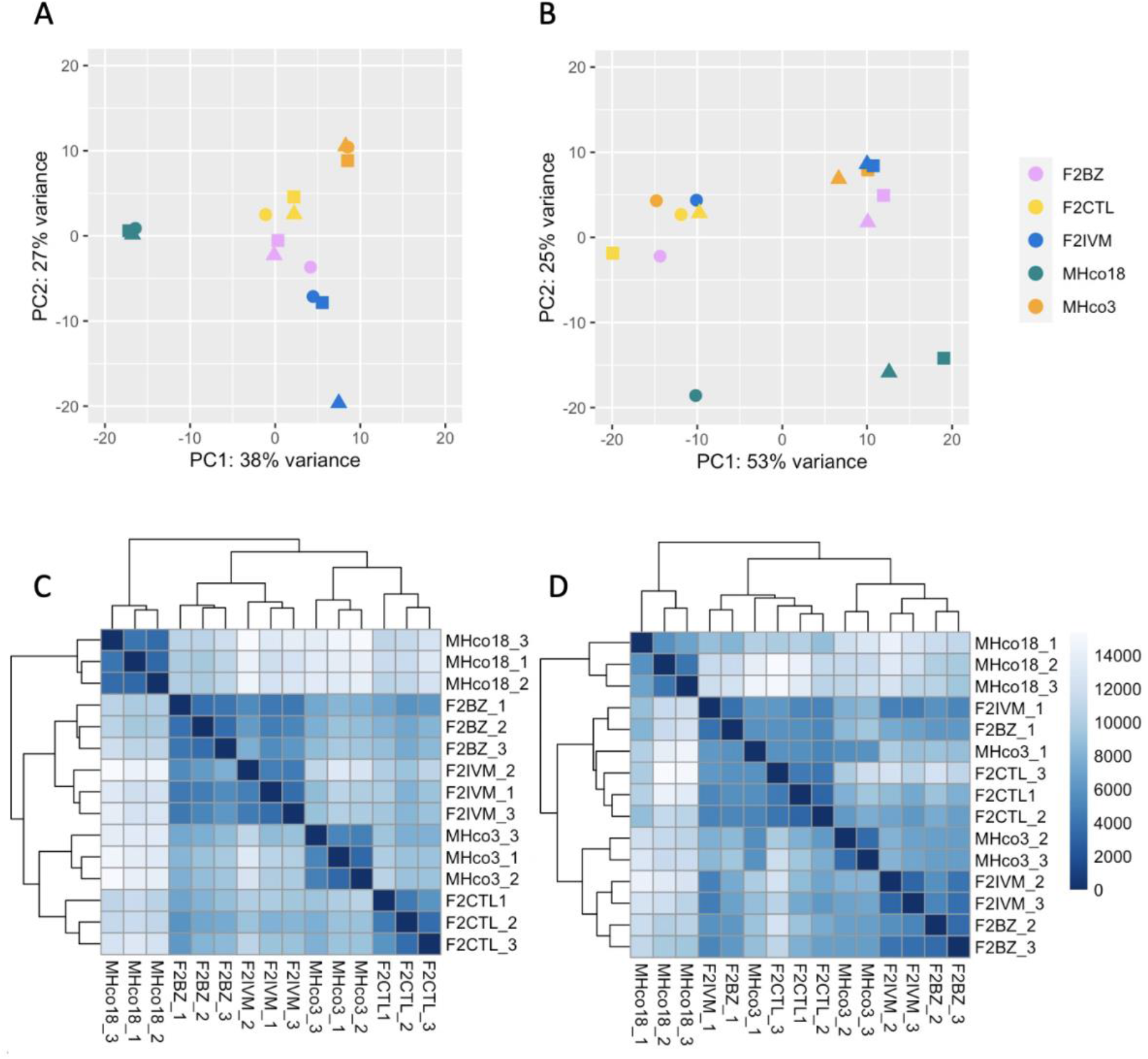
PCA plots (A. males, B. females) with regularised log transformed data, based on the 500 most variable genes, and heatmaps (C. males, D. females) showing Poisson distance between samples. MHco3 = drug-susceptible parent, MHco18 = drug-resistant parent, F2IVM = ivermectin-treated genetic cross F2, F2BZ = benzimidazole-treated F2, F2CTL = untreated control F2. The shape of point in the PCA and the sample number in heatmap relate to the order that the samples were processed (circle = 1st, triangle = 2nd, square = 3rd).

### Separating transcriptomic differences associated with drug resistance from other between-isolate variation

To investigate whether ivermectin resistance was associated with differential gene expression, we undertook multiple pairwise comparisons (Figure S1, Figure S4). We predicted that the MHco3 and MHco18 isolates would have many constitutive differences in gene expression reflecting genetic diversity resulting from their distinct evolutionary histories, including differences in their sensitivity to the benzimidazoles, levamisole and ivermectin. However, we predicted that strain-specific and drug-specific differences could be separated using the genetic cross: the subset of genes specifically associated with resistance to ivermectin would be differentially expressed in the ivermectin selected genetic cross relative to the unselected genetic cross, but strain-specific differences would not. Consistent with these expectations, the largest number of differentially expressed genes that were unique to a pairwise comparison was between parental isolates (n = 1481 and 1290, males and females respectively, Figure S4). Genes that were only differentially expressed in the parental isolates (Figure 2, grey points) were essentially ruled out as playing a role in ivermectin resistance mediated by gene expression because they were not also differentially expressed in the ivermectin selected genetic cross (Figure 2, blue points), with the caveat that our analysis was by choice conservative being based on a stringent cut off (adj P < 0.01) for every pairwise comparison. Using this approach, we identified differentially expressed genes that were specifically associated with ivermectin resistance.

**Figure 2.**
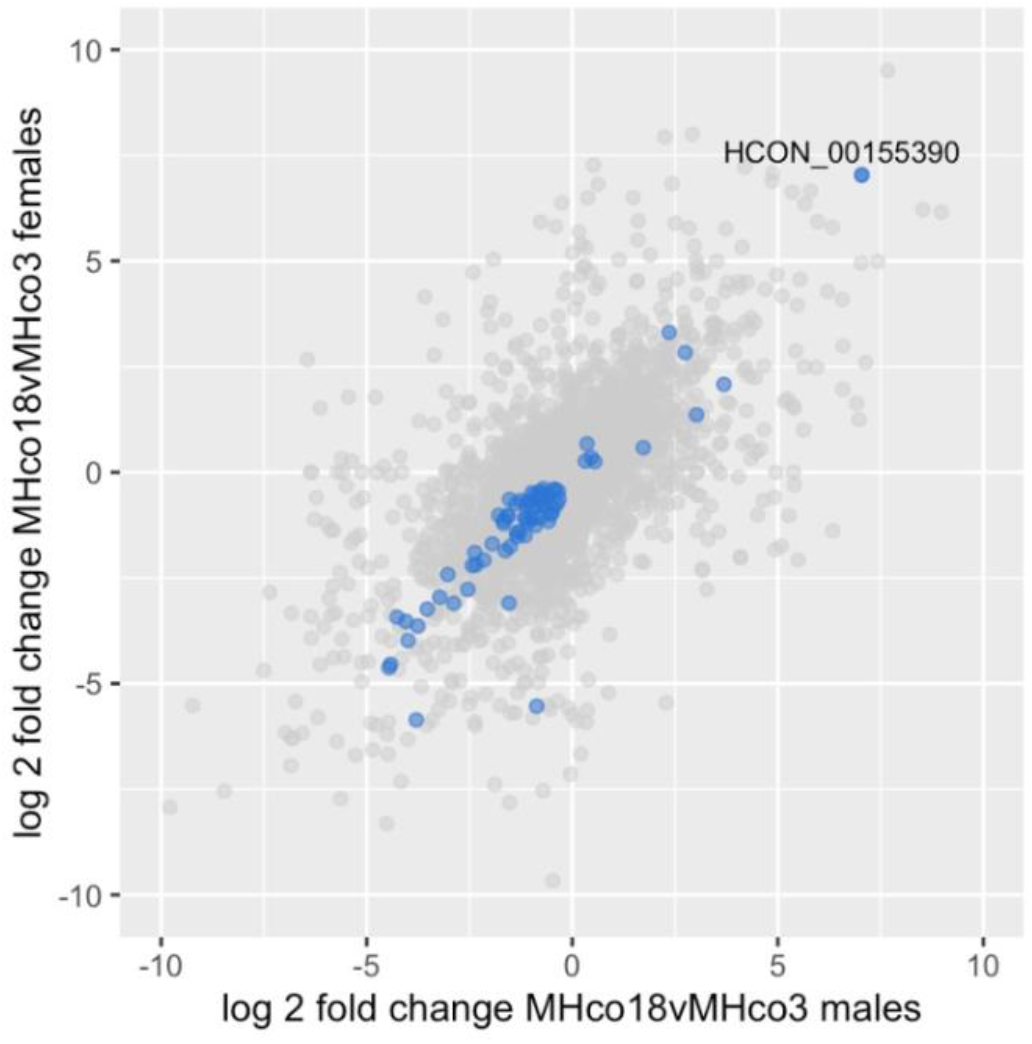
Scatter plot showing differentially expressed genes in comparisons of resistant and susceptible populations (adj P < 0.01), with log2 fold change of males shown on the x-axis and females on the y-axis. Grey points represent genes that are differentially expressed only in parental (MHco18 vs MHco3) comparisons, i.e. likely to represent between-strain differences not associated with ivermectin resistance. If blue they are also differentially expressed in the F2IVM vs MHco3 and F2IVM vs F2CTL comparisons, i.e. likely to be specifically associated with ivermectin resistance. The most highly upregulated gene in resistant isolates is the *H. contortus* homologue of *C. elegans cky-1* (HCON_00155390).

### Differentially expressed genes are enriched within the ivermectin QTL on chromosome V after ivermectin selection in the genetic cross

We have previously identified a major effect QTL on chromosome V in response to ivermectin selection in laboratory strains and in field samples of *H. contortus* [17, 18]. We inspected the distribution of differentially expressed genes at this locus and throughout the genome for each pairwise comparison (Figure S5). These plots highlighted the large differences in gene expression between parental isolates (S5, A and B) and a reduction in the degree of transcriptomic differentiation in comparisons using the ivermectin treated genetic cross (S5, C, D and E). In comparisons between parental isolates, differentially expressed genes were present throughout the genome with no enrichment for differential expression in the ivermectin resistance locus centred around 37.5 Mb on chromosome V (*X*^2^ = 2.62, P = 0.11). However, the subset of these genes that were also differentially expressed in all comparisons with the genetic cross post ivermectin selection (F2IVM) showed significant enrichment for genes located in the ivermectin resistance locus (*X*^2^ = 37.76, P = 7.23E-10).

### Upregulation of a single gene within the ivermectin QTL in all resistant isolates

Seventy-six genes were up- or downregulated in all male and female ivermectin resistant populations (Figure S1B, Figure 2, Table S1). Sixty of the differentially expressed genes had homologues in *C. elegans* (Table S1) but this gene set showed no tissue, phenotype or GO term enrichment from *C. elegans* data. The gene with highest upregulation, *HCON_00155390*, a homologue of *C. elegans cky-1*, lies within the major chromosome V locus under ivermectin selection in *H. contortus* (Figure 3). As shown in Table S1, *HCON_00155390:cky-1* is expressed between 62 and 132-fold higher (log2 fold changes from 5.95 to 7.04, adj P ≤ 0.00024) in males and females of all resistant populations relative to the susceptible MHco3 isolate and 2.7-fold higher (1.44 log2 fold change, adj P = 0.0019) in males of the genetic cross after ivermectin selection (F2IVM = resistant genotypes only) relative to the unselected population (F2CTL = mix of resistant and susceptible genotypes). Constitutive upregulation of *HCON_00155390:cky-1* was confirmed in MHco18 males relative to MHco3 males with RT-qPCR [18]. Constitutive upregulation in MHco18 females was also detected relative to MHco3 females, but the fold-change in expression could not be calculated due to a lack of measurable expression in MHco3 females (data not shown). In *C. elegans, cky-1* encodes a transcription factor expressed in the embryo and in non-neuronal pharyngeal cells [19] and the only published phenotype is early onset of polyglutamine-YFP aggregation after RNAi [20]. The human orthologue, *NPAS4* or *NXF*, is a basic helix-loop-helix transcription factor that regulates gamma-aminobutyric acid (GABA) releasing inhibitory synapses [21] and can be induced in response to neurodegeneration or excitation to confer protection to neuronal cells [22].

**Figure 3.**
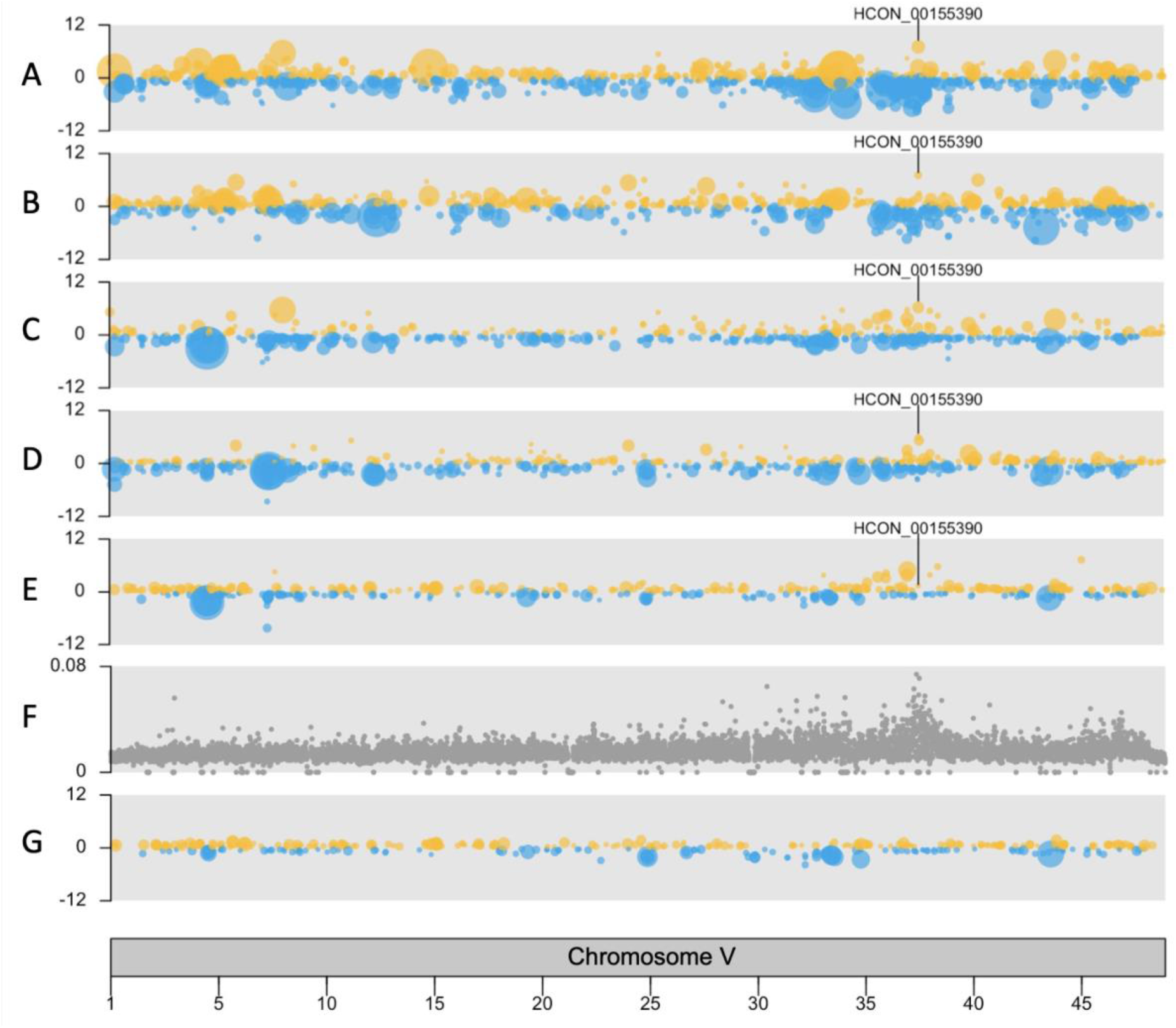
Chromosome V karyoplots showing genomic position of genes with significant upregulation (yellow) or downregulation (blue). Each point represents a differentially expressed gene (adj P < 0.01) and point size corresponds to significance. Panels A-D compare ivermectin-resistant versus ivermectin-susceptible populations (A. MHco18 vs MHco3 males, B. MHco18 vs MHco3 females, C. F2IVM vs MHco3 males, D. F2IVM vs MHco3 females). Panel E compares an ivermectin-resistant versus a mixed (resistant and susceptible) population (F2IVM vs F2CTL males). Panel F shows genetic differentiation (*F*_ST_) between the F3 generation of the genetic cross pre- and post-ivermectin selection to highlight the major locus of ivermectin selection [18]. Panel G shows relative absence of differential expression at the ivermectin QTL in a comparison with a benzimidazole selected population (F2BZ vs F2CTL males). Upregulation of the *H. contortus* homologue of *C. elegans cky-1* (HCON_00155390) is identified in ivermectin resistant populations only.

One of the most highly downregulated genes also lies within the major locus under ivermectin selection: *HCON_00155240*, a homologue of *C. elegans* F59B1.10. In *C. elegans*, this gene encodes a checkpoint (CHK) kinase-like protein with an uncharacterised oxidoreductase Dhs-27 domain, expressed in GABAnergic neurons, dopaminergic neurons and muscle cells. One caveat to our measurement of expression for this gene is its high level of polymorphism in the MHco18 isolate, which could give the appearance of downregulation due to a greater proportion of unmapped reads relative to MHco3 (discussed in more detail below). Outside of the ivermectin QTL, downregulated genes included seven homologues of *C. elegans vap-1*, a cysteine rich secretory protein expressed in the amphid sheath cell.

Notably, no putative ivermectin target or resistance genes from the literature [7, 8, 11, 23, 24] were identified as differentially expressed in all resistant populations; normalised read counts for candidate genes are shown in Figures S6 (males) and S7 (females). Genes that are differentially expressed in the parental isolates only are unlikely to be related to ivermectin resistance and none of these candidate genes lie within genomic loci under ivermectin selection [17, 18].

### Male and female worms show shared and sex-specific differences in gene expression associated with ivermectin resistance

We were concerned that the higher between-replicate variability of the female data could limit the detection of truly differentially expressed genes in the male analysis. It is also possible that ivermectin affects male and female worms differently [25, 26]. For these reasons, we also analysed results from the male and female datasets separately (Figure S1B, Figure 4). In the male worms, there were an additional 67 upregulated and 107 downregulated genes that were not differentially expressed in all pairwise comparisons of ivermectin resistant female worms (Table S2). There was low but significant upregulation of two putative multi-drug resistance proteins associated with ivermectin resistance in the literature: *HCON_00162780* and *HCON_00175410*, the homologues of *C. elegans pgp-11* and *mrp-4*, respectively. PGP-11 is a P-glycoprotein known to modulate ivermectin sensitivity in other nematode species [27], and the *pgp-11* locus has previously been associated with ivermectin resistance in *H. contortus* [16, 17, 24]. In this study, the expression of *HCON_00162780:pgp-11* was markedly higher in males relative to females (Figure 5A and 5B), and upregulation of *pgp-11* in MHco18 versus MHco3 males was confirmed with RT-qPCR (Figure 5C). In *C. elegans, mrp-4* is involved in lipid transport and lipid storage, but has also been shown to have higher constitutive expression in an ivermectin resistant strain of *C. elegans* relative to wild-type [28]. One putative ligand gated ion channel, *HCON_00137455*, a homologue of *lgc-25*, was downregulated, as was *HCON_00078660*, a homologue of the sodium leak ion channel *nca-2*. In *C. elegans, nca-2* is expressed in acetylcholine and GABA motor neurons [29], and is involved in the propagation of neuronal activity from cell bodies to synapses [30] and in synaptic vesicle recycling [29]. There was upregulation of one chloride channel, *HCON_00108990*, the homologue of *best-19*. In *C. elegans*, this is a bestrophin chloride channel expressed in the NSM, a pharyngeal neurosecretory motor neuron.

**Figure 4.**
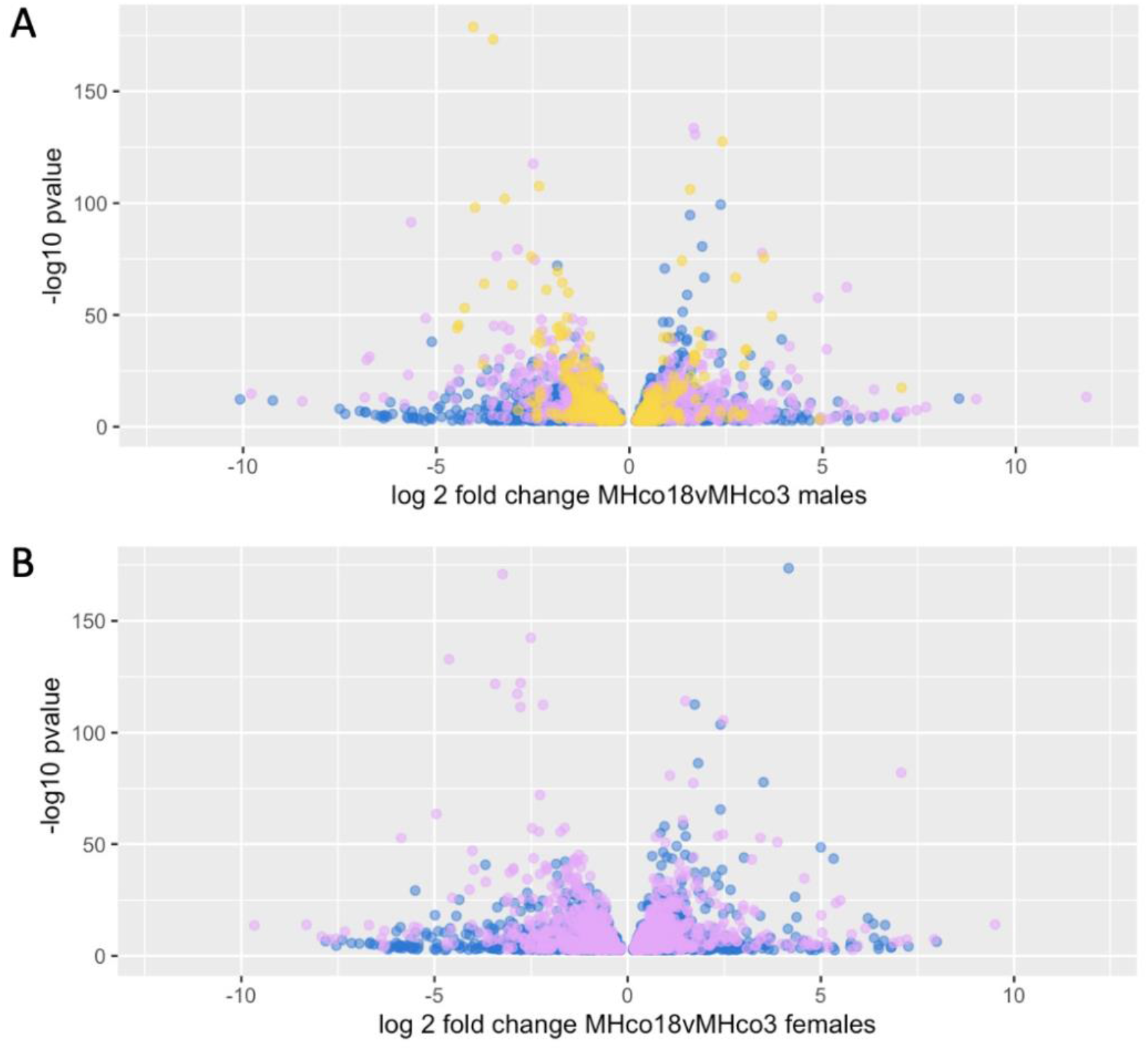
Volcano plots showing differentially expressed genes in male (A) and female (B) samples (adj P < 0.01). Blue points represent genes that are only differentially expressed in the parental (MHco18 vs MHco3) comparisons. If purple, they are also differentially expressed in the F2IVM vs MHco3 comparisons. If yellow (males only), they are also differentially expressed in the F2IVM vs F2CTL comparison.

**Figure 5.**
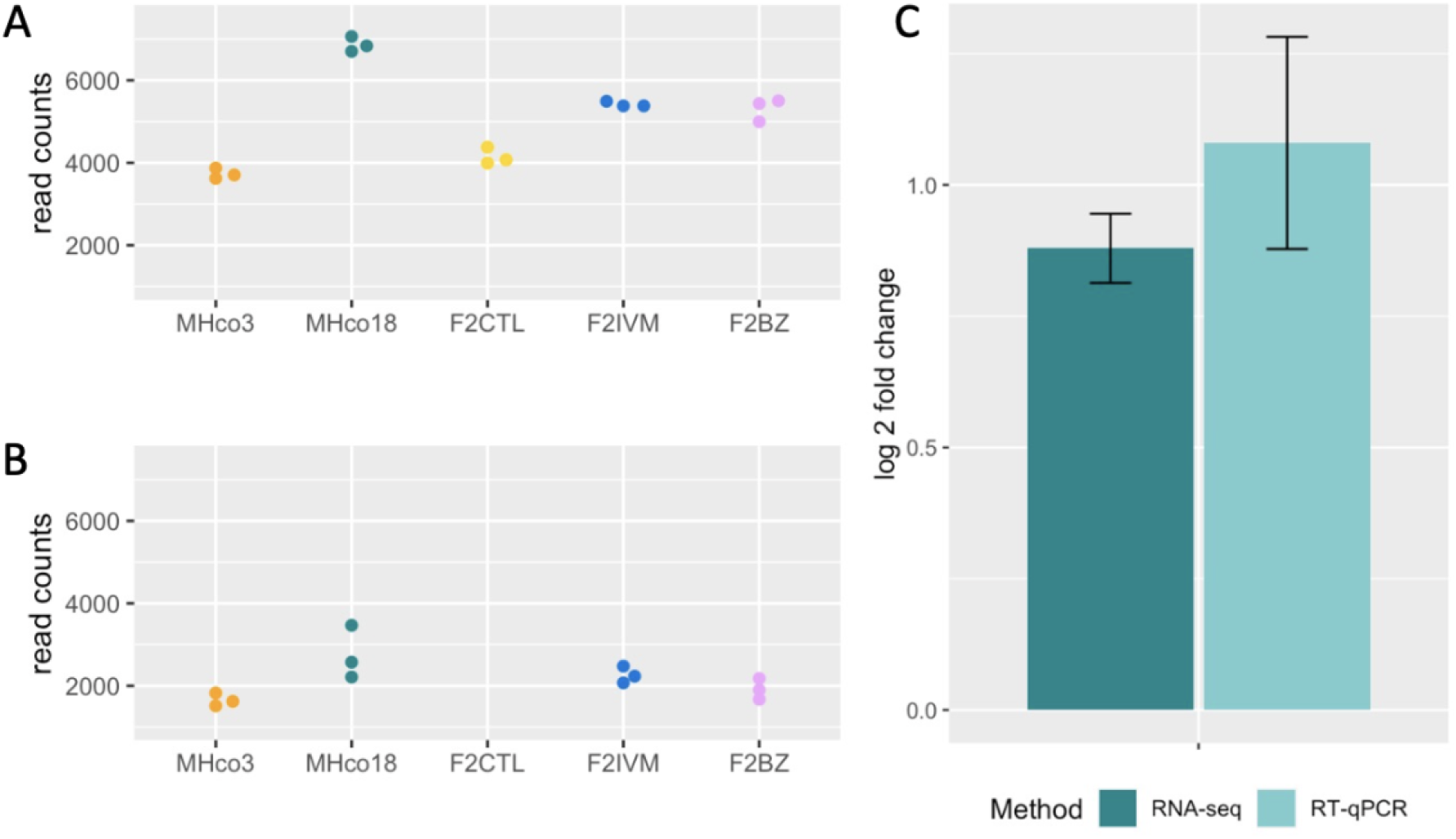
Expression analysis of *HCON_00162780:pgp-11*. Normalised read counts for male (A) and female (B) RNA-seq samples highlighting higher constitutive expression in all male populations relative to female populations. C. Comparison of RNA-seq and RT-qPCR data (mean of three biological replicates) for MHco18 males relative to MHco3 males, showing constitutive upregulation in the ivermectin resistant isolate. For RNA-seq data, error bars show the log fold change standard error (lfcSE) and for RT-qPCRs, error bars show the standard deviation.

Of the differentially expressed genes in ivermectin resistant males, 125 had *C. elegans* homologues (Table S2). These homologues were enriched for expression in the ALA neuron (E = 0.24, O = 4, Q = 0.0011) (Figure S8), a mechanosensory interneuron that projects into the nerve ring then joins the lateral cords, extending as far as the tail, adjacent to the excretory canals [31]. The ALA neuron inhibits locomotion and pharyngeal pumping during lethargus and ‘stress induced sleep’, a characteristic behaviour in *C. elegans* during recovery from exposure to damaging conditions [32, 33]. There was GO enrichment for a single term ‘neuropeptide signalling pathway’ (E = 0.92, O = 6, Q = 0.013), associated with *nlp-9, nlp-14, flp-6, flp-7, flp-16* and *flp-24*, all of which were downregulated. Notably, there was also phenotype enrichment for 13 terms associated with movement (Figure S8).

For female worms, the lack of an F2CTL sample meant the equivalent analysis (excluding differentially expressed genes in all resistant male comparisons) would be very likely to be confounded by differences between the female parental isolates that were not associated with ivermectin resistance (Figure 4B). To mitigate this, an alternative analysis including previously published RNA-seq data for females from two additional ivermectin resistant populations (MHco4(WRS) and MHco10(CAVR) versus MHco3 [34]) was undertaken (Figure S1B). This identified 85 upregulated and 97 downregulated genes in all ivermectin resistant female samples that were not differentially expressed in ivermectin resistant male samples (Table S3). Downregulation of *HCON_00076370*, a homologue of the *C. elegans* sodium:potassium:chloride symporter *nkcc-1*, was identified in this analysis. In *C. elegans, nkcc-1* is a chloride transporter and functions as a developmental switch that changes the response of GABA_A_ receptors in body wall muscles from depolarising in the early L1 to hyperpolarising from the mid-L1 stage onwards by controlling the cellular chloride gradient [35]. In female worms only, there was also downregulation of one putative ligand gated ion channel, *HCON_00097440*, a homologue of *acr-24*. Of the differentially expressed genes in ivermectin resistant females, 158 had *C. elegans* homologues. These showed no tissue or GO term enrichment but there was enrichment for a single phenotype ‘uptake by intestinal cell defective’ (E = 0.59, O = 4, Q = 0.075).

### Transcriptomic response to ivermectin exposure implicates neuronal regeneration, neuropeptide signalling and chloride homeostasis

Having identified genes with constitutive differential expression in ivermectin resistant worms, we were also interested in genes that were only up or downregulated after exposure to ivermectin treatment. This subset of genes would be expected to be differentially expressed in the F2IVM versus F2CTL comparison and the F2IVM versus MHco3 comparisons, but not the MHco18 versus MHco3 comparisons, because the MHco18 isolate had not been exposed to ivermectin. As ivermectin has a short half-life in sheep (plasma t_1/2_ = 2.7 days [36]) and the F2IVM worms were isolated 21 days after ivermectin treatment, this gene set will not reflect acute ivermectin exposure but rather the longer-term influence of drug treatment.

Of 120 genes that were differentially expressed in response to ivermectin treatment, 95 had *C. elegans* homologues (Table S4). The most highly upregulated gene with a *C. elegans* homologue was *HCON_00068595* which was 5 to 32-fold more highly expressed in ivermectin treated populations (2.31 – 4.99 log2 fold change, adj P ≤ 2.52E-12 (Table S4)). This gene is a homologue of *emb-9*, a collagen type IV alpha 5 chain, involved in many processes such as pharyngeal pumping and egg laying, but with a critical role in positive regulation of neuron regeneration following axon damage [37]. *emb-9* is most strongly expressed in the basement membrane of the *C. elegans* pharynx [38].

There was no differential expression of any ligand gated ion channels after ivermectin exposure, but *HCON_00068600*, the homologue of a neuronal CLC-2 chloride channel, *clh-3*, was upregulated between 2 and 12-fold (1.28 – 3.62 log2 fold change, P ≤ 2.95E-10 (Table S4)) in ivermectin treated populations. CLC-2 is activated by membrane hyperpolarisation and is thought to regulate neuronal excitability by providing an efflux pathway for chloride. In *C. elegans, clh-3* is only expressed in the (spontaneously active) hermaphrodite-specific motor neuron (HSN), where *clh-3* inhibits excitability by mediating chloride influx, resulting in suppression of egg laying [39]. In *H. contortus, HCON_00068600:clh-3* is also constitutively expressed in both male parental samples, suggesting expression is not limited to the HSN in this species. Interestingly, HCON_00068595:*emb-9* and HCON_00068600:*clh-3* are neighbouring genes on chromosome III, although encoded on opposite strands. In *C. elegans, emb-9* and *clh-3* are encoded on different chromosomes (chromosome III and II, respectively) and have no known interaction.

For the *C. elegans* homologues, the most significant GO term was ‘neuropeptide signalling pathway’ (E = 0.71, O = 8, Q = 6e-06) (Figure S8). The eight genes associated with this term encoded five neuropeptide-like proteins (*nlp-7, nlp-11, nlp-13, nlp-49* and *nlp-50*) and three FMRF-like peptides (*flp-1, flp-14* and *flp-22*); all were downregulated in F2IVM samples. In common with the constitutive datasets, multiple homologues of *C. elegans vap-1* (15 in total) were downregulated.

### Differential expression of genes associated with neuronal plasticity and chloride homeostasis are specific to ivermectin selection

To investigate the specificity of the transcriptomic differences we identified in the ivermectin-selected genetic cross and to explore the possibility of a common response to drug exposure (for example, a stress response or multi-drug resistance pathway), we also undertook RNA-seq analysis of the same F2 population after benzimidazole selection. There were fewer differentially expressed genes in the benzimidazole resistant populations relative to the equivalent comparisons for ivermectin (59 associated with benzimidazole resistance (Table S5) versus 76 for ivermectin, and 55 in response to benzimidazole treatment (Table S6) versus 120 for ivermectin). The benzimidazole resistant populations showed no differential expression of genes known to confer resistance through non-synonymous mutations in the target protein, β-tubulin (i.e., *β-tubulin isotype 1* (*HCON_00005260*) or *β-tubulin isotype 2* (*HCON_00043670*)) or genes associated with drug metabolism (UDP-glycotransferases or cytochrome P450 enzymes).

Notably, both the ivermectin and benzimidazole-selected populations appeared to be transcriptionally very similar (Figure 1), despite different regions of the genome being under selection by each drug, with no evidence of a shared genetic component to resistance [18]. In the benzimidazole resistant and response to treatment groups, 35 genes in each (59% and 65% of differentially expressed genes, respectively) were shared with the equivalent ivermectin selected populations. In the benzimidazole resistant populations, as for the ivermectin resistant populations, there was downregulation of multiple homologues of *C. elegans vap-1* (five downregulated constitutively and 11 in response to benzimidazole treatment) and four of the same neuropeptides were downregulated in response to benzimidazole and ivermectin treatment (homologues of *nlp-13, flp-1, flp-14* and *flp-22*). In male worms only, upregulation of one putative multi-drug resistant gene, *HCON_00162780:pgp-11*, was identified in all pairwise comparisons for both ivermectin and benzimidazole resistant populations.

However, in contrast with the ivermectin resistant populations, there was no enrichment for differential expression at the chromosome V locus (Figure 3, bottom panel) and no differential expression of *HCON_00155390:cky-1* or any of the genes associated with neuronal regeneration and chloride homeostasis. This suggests the differential expression of these genes, identified using the same genetic cross, are unique to ivermectin resistance and recovery.

### Use of a genetic cross minimises the impact of between-isolate polymorphism on differential expression analysis

Previous work highlighted the potential impact of high levels of polymorphism on between-isolate RNA-seq analysis in *H. contortus* [34], showing that polymorphic isolates (relative to the reference MHco3 isolate) had significantly fewer reads mapping to the genome, potentially biasing differential expression analysis. In this study, the difference in ‘SNP rate’ (the number of SNPs in a gene model; see Methods) between parental isolates had a small but significant negative correlation with gene expression (R^2^ = 0.074 [P < 0.001] and R^2^ = 0.099 [P < 0.001] for males and females, respectively). This correlation suggests that highly polymorphic genes were more likely to be identified as downregulated than upregulated in the parental isolates (Figure S9). This effect was more striking for the MHco18 isolate (upper left quadrant) but could be seen to a lesser extent for polymorphic genes in the MHco3 isolate (lower right quadrant). Admixture of the parental genotypes in the genetic cross largely mitigated this bias in male analyses (R^2^ = 0.0026 [P = 0.322]) (Figure S9, dark blue points), although the lack of a female F2CTL limited our ability to control for polymorphism in the female-only analyses (R^2^ = 0.074 [P < 0.001]).

The positive impact of the cross in the male analyses was further supported by a reduction in the proportion of highly polymorphic genes (defined as having a SNP rate of greater than 2% in MHco18) in the downregulated genes identified in the F2IVM versus F2CTL analysis compared to the MHco18 versus MHco3 analysis. Notably, 21% (469/2196) of genes identified as downregulated in MHco18 males were highly polymorphic, relative to 2.8% (58/2069) of upregulated genes, whereas in the genetic cross 6.2% (60/973) of genes identified as downregulated in F2IVM males were highly polymorphic, relative to 5.8% (71/1219) of upregulated genes. Finally, the relevance of controlling for polymorphism was highlighted by the subset of genes that were identified as the most highly polymorphic in the parental isolates: *C. elegans* homologues of the 1684 genes with a greater than 2% difference in polymorphism rate in the MHco18 isolate relative to the MHco3, were enriched for the phenotype ‘anthelmintic response variant’ (E = 3.2, O = 10, Q = 0.058). This was associated with highly polymorphic homologues of *avr-15, acr-16, lev-1, osm-1, che-1, che-3, che-11, che-14, daf-10* and *dyf-3*. Although not identified in the enrichment analysis, a homologue of *ben-1, HCON_00043670* (β-tubulin isotype 2), was also highly polymorphic. Overall, these findings highlight the need to control for polymorphism in transcriptomic analyses of genetically divergent populations and find that genes associated with drug response are among the most polymorphic genes in a resistant population.

## Discussion

A key challenge of genomic analyses of helminths is accounting for the high genetic diversity between populations [40]. Comparisons between resistant and susceptible populations are further complicated by inherent differences in traits other than resistance [16]. In the case of transcriptomic analyses, the same factors confound detection of differential gene expression specifically associated with resistance, and commonly used measurements of gene expression can be biased by between-population polymorphism [34]. Further, for transcriptomic analyses, differential gene expression after drug selection may reflect constitutive differences in gene expression between resistant and susceptible worms and/or differences that are induced by drug exposure. The latter may be specific to a particular drug or may represent a more generic stress response or drug metabolism pathway. Separating these different mechanisms in between-population comparisons is complex and usually not feasible.

In this study, we used a genetic cross of a multi-drug resistant isolate of *H. contortus* with a susceptible (reference) isolate, providing a population composed of an admixture of parental genotypes, to control for differences in the genetic background of each parent [18] and to mitigate the impact of polymorphism on downstream analyses. Comparisons of resistant adults with and without ivermectin selection allowed us to separate differential expression associated with ivermectin resistance from that associated with the response to drug exposure. Further, a comparison of genes with differential expression after selection with ivermectin or benzimidazole allowed us to identify drug-specific and shared responses.

The success of the genetic crossing approach in reducing the impact of background genetic diversity between the parental isolates was demonstrated by the reduction in the number of differentially expressed genes identified in the genetic cross with and without ivermectin treatment, relative to the number of differentially expressed genes identified between parents. In the genetic cross, there was also a reduction in the distribution of differentially expressed genes throughout the genome, with significant enrichment for differentially expressed genes within the ivermectin resistance QTL on chromosome V, which was not detectable in the parental comparison due to high transcriptomic diversity throughout the genome.

A previous study highlighted the potential confounding effect of genetic diversity on RNA-seq analysis of different *H. contortus* isolates due to poorer mapping of divergent reads to the reference genome [34]. In this study, alignment of longer reads (150 bp vs 100 bp) to a more contiguous genome assembly [14] with an improved aligner (HISAT2 [41]) resulted in more reads from the divergent MHco18 isolate aligned to the reference genome (75-77%), than from the reference isolate in the previous study (69%) [34]. However, the small but significant negative correlation between level of polymorphism and differential expression in the parental isolates suggests that mapping biases could still impact the accuracy of differential expression measurement for polymorphic genes in between-isolate comparisons. Further, the significant enrichment for an ‘anthelmintic response variant’ phenotype in the most highly polymorphic subset of genes highlights a challenge of studying complex traits in rapidly evolving helminths: the polymorphic genes may be those of most interest, but their analysis can be confounded when relying on a single reference genome for read alignment and/or primer design, or when comparing sequences from different populations to infer a relationship with the trait of interest. Improved tools for mapping RNA-seq reads to a reference genome, such as HISAT-genotype [42], which incorporates known variants to optimise the mapping of polymorphic regions of the human genome is one promising approach, although such an approach is not yet available for non-model organisms. However, in this study, the use of the genetic cross resulting in an admixture of parental genotypes largely corrected for artefactual differential expression due to mapping bias in male samples, although some bias remained for females due to the lack of a F2 control.

Notably, after controlling for differences in the genetic background of our resistant and susceptible isolates with the genetic cross, we found no evidence of differential expression of candidate ivermectin resistance genes from the literature [7, 8, 11, 23, 24], with the exception of *HCON_00162780:pgp-11*, which was upregulated in resistant males only. While this does not rule out a role for differential expression of these genes in the early stages of resistance development, it suggests they do not confer high level resistance; this is consistent with our genomic analyses of ivermectin resistance, where we find no candidate resistance genes in the major locus under ivermectin selection [17, 18]. Despite finding no evidence of differential expression of any glutamate gated chloride channel receptor subunits, the predicted major targets of ivermectin [43-45], we did identify differential expression of multiple neuronal genes predicted to modulate cellular chloride levels (*clh-3, best-19, nkcc-1*), which could potentially mitigate the effects of chloride influx when ivermectin binds to its receptor.

The gene with the most robust association with ivermectin resistance was the homologue of *C. elegans cky-1*, a gene with no previous association with drug resistance. *HCON_00155390:cky-1* lies at the centre of the major locus of ivermectin resistance on chromosome V [18] and was the most highly upregulated gene overall in both males and females. In *C. elegans, cky-1* is a transcription factor encoding basic helix-loop-helix (bHLH) and Per-Arnt-Sim (PAS) domains; these domains are also present in the *H. contortus* homologue. Very little is known about the function of *cky-1* in nematodes, but the mammalian orthologue (NPAS4/NXF) regulates GABA releasing inhibitory synapses [21] and is induced by neurodegeneration or excitation to confer protection to neuronal cells [22]. There are many hundred predicted targets of NPAS4 in the mouse, based on differential expression measured by microarray after NPAS4-RNAi [21], including various ion channels and synaptic proteins, but many (more than a third) of the target genes are uncharacterised. In *C. elegans*, only four *cky-1*-interacting genes are listed in WormBase: *aha-1, pmk-1, vab-3* and *ztf-2*. Homologues of all four genes are present in the *H. contortus* genome but none are differentially expressed in all ivermectin resistant adults. It is possible that some of the other differentially expressed genes identified in this study are downstream targets of *HCON_00155390:cky-1* and this warrants further investigation. In *C. elegans, cky-1* expression is highest in the embryo and is largely restricted to pharyngeal cells (arcade cells, pharyngeal muscle cells, hypodermis, and pharyngeal:intestinal valve) with little or no expression detected elsewhere by single cell RNA-seq [19]. It is unknown if this pharyngeal expression pattern is the same in either ivermectin susceptible or resistant *H. contortus* but studies are underway. The major routes of ivermectin uptake in adult *H. contortus* are unclear, but it is likely that the pharynx plays a key role.

Despite surviving treatment, resistant nematodes can show phenotypic effects after ivermectin exposure, for example, reduced pumping in the triple resistant *C. elegans* DA1316 [46] and suppression of egg production in gastrointestinal nematodes [47], particularly at the early stages of resistance development, which suggests they are still physiologically affected to some degree by the drug. These phenotypic effects could relate to the many potential targets of ivermectin [48] and/or the mode of resistance. We identified significant enrichment for downregulation of multiple inhibitory neuropeptides after drug exposure (and with constitutive differential expression in resistant males). These signalling peptides regulate various aspects of behaviour, including locomotion, pharyngeal pumping, lipid storage and egg laying, which may relate to phenotypes observed in ivermectin treated worms. Transcriptomic data also highlighted genes associated with neuronal plasticity, neuronal development and regeneration, yet many of these genes are pleiotropic, and differential expression may equally relate to their roles in overcoming the phenotypic effects of ivermectin on feeding, movement and reproduction. For example, in *C. elegans, emb-9* positively regulates neuronal regeneration following damage yet is also essential for pharyngeal pumping and egg laying [37].

Our analysis of the same genetic cross after benzimidazole selection supports our expectation that the genes we have identified with putative roles in ivermectin resistance are largely unique to this drug. However, the degree of overlap in the two groups was higher than expected given the separate modes of action and distinct genomic loci under selection by each drug [18]. One technical explanation could be the impact of the male F2CTL sample: if, as seen in the female data, this sample was an outlier, it could artificially inflate the number of shared differentially expressed genes in the F2BZ and F2IVM samples. Despite passing all our QC analysis, there was some evidence to support this in the higher number of genes that were differentially expressed in the male F2IVM versus F2CTL comparison relative to the male F2IVM versus MHco3 comparison, which is hard to explain biologically. However, even if this is the case, based on the PCA plots, the benzimidazole-selected and ivermectin-selected populations do appear to be transcriptomically similar. Another explanation would be stochastic inheritance of the same parental alleles at regions of the genome that are not necessarily under selection by both drugs; this would reflect the divergence of the two parental isolates and is increasingly likely with a small sample size (in this case, 20 worms per sample), in a polyandrous species [49], and with a limited number of recombination events (F2 generation of the cross). Alternatively, the results could indicate a degree of co-selection of mechanisms promoting tolerance of multiple anthelmintics in a population routinely treated with all drug classes. However, other than *pgp-11*, which may represent a multi-drug resistant protein, there were no known resistance-associated genes or pathways shared between the ivermectin and benzimidazole selected groups.

One particularly striking finding was the apparent lack of differential expression of genes involved in xenobiotic metabolism for either drug class; this is consistent with the expectation that ivermectin is not metabolised in nematodes [46], but is more surprising for the benzimidazoles where biotransformation has been detected in *H. contortus* and *C. elegans* [50-52]. This may reflect the extended time between treatment and isolation of worms in this study, which would miss an acute response to circulating drug. However, our results are consistent with a recent study investigating the acute transcriptional response of adult MHco18 *H. contortus* exposed to different benzimidazoles *in vitro*, where, in contrast to the equivalent experiment in *C. elegans*, no genes with roles in xenobiotic metabolism were differentially expressed [53].

In conclusion, the use of a genetic cross has allowed the separation of transcriptomic differences associated with ivermectin resistance and survival from a background of high transcriptomic differentiation in resistant and susceptible isolates of *H. contortus*. We identify constitutive upregulation of *HCON_00155390:cky-1*, a gene in the centre of the major genomic locus under ivermectin selection in global populations of *H. contortus*. We also identify a small number of differentially expressed genes outwith the genomic locus, which may be downstream targets of *cky-1* or may function in shared or parallel pathways to promote ivermectin resistance.

## Materials and Methods

### Generation of parasite material

Fifteen donor sheep were treated with 2.5 mg/kg bodyweight monepantel (Zolvix Oral Solution for Sheep, Elanco AH) 14 days prior to infection to ensure they were parasite free. Each donor was orally infected with 5000 L_3_ of a single *H. contortus* isolate; three donors were infected with the drug susceptible MHco3(ISE) isolate [54], three donors were infected with the triple resistant MHco18(UGA2004) isolate (benzimidazole, levamisole and ivermectin resistant) [7] and nine donors were infected with the F2 generation of a genetic cross between MHco3 females and MHco18 males [15]. For the genetic cross F2 infections, on day 35 post-infection (pi), three donor sheep were treated with 0.2 mg/kg body weight ivermectin (Oramec Drench, Boehringer Ingelheim Animal Health UK Ltd) (F2IVM), three were treated with 7.5 mg/kg fenbendazole (Panacur 10% Oral Suspension, MSD Animal Health) (F2BZ), and three were left untreated (F2CTL). Adult worms were harvested at post mortem, sexed and snap frozen in liquid nitrogen for storage at -80°C in batches of 20 worms. Post mortem dates differed for each isolate: day 62 pi for MHco3, day 28 pi for MHco18 and day 56 pi for the F2 infections.

### RNA isolation and library preparation for RNA-seq

Adult worms from one donor each for the MHco3, MHco18, F2IVM, F2BZ and F2CTL populations were used for RNA-seq. Total RNA was isolated from three batches of 20 male or female worms per donor using a standard Trizol (Thermo Fisher Scientific, 15596026) extraction. Each RNA sample underwent DNaseI (Qiagen, 79254) treatment in solution followed by purification on a RNeasy Mini Spin Column (Qiagen, 74104), before freezing at -80°C. Total RNA (1 μg) from each sample was used for NEBNext PolyA selection and Ultra Directional RNA Library preparation (Illumina, E7490, E7420). Prepared libraries were sequenced using 2×150 bp paired-end read chemistry on an Illumina HiSeq 4000 platform.

### Messenger RNA data processing

Raw Fastq files were trimmed for the presence of adapters using Cutadapt v1.2.1 [55]. Option -O 3 was used to trim the 3’ end of any reads matching adapter sequence for 3 bp or more. The reads were further trimmed using Sickle v1.200 [56] with a minimum window quality score of 20. Reads shorter than 20 bp after trimming were removed. Downstream analyses were undertaken in Galaxy [57] and R Studio v1.1.456 [58]. Sequence quality was assessed with FastQC [59] and MultiQC [60]. Reads were aligned to the *H. contortus* V4 genome (haemonchus_contortus_MHCO3ISE_4.0 [14]) with HISAT2 v2.1.0 using default settings [42]. Counts of reads mapping to gene models (haemonchus_contortus.PREJEB506.WBPS15.annotations.gff3) were determined with *featureCounts* in Subread v1.6.2 [61] with the -O option to allow reads to contribute to multiple features. Differential expression analysis was undertaken with DESeq2 v1.22.2 [62] with alpha set to 0.01.

### Genomic DNA data processing

The ‘SNP rate’ for every gene was calculated from whole genome sequencing data from populations of 200 L_3_ (Pool-seq) from the MHco3 and MHco18 isolates [18] as described previously [34]. Briefly, biallelic SNPs relative to the *H. contortus* V4 genome (haemonchus_contortus_MHCO3ISE_4.0 [14]) with allele frequency >0.4 within the population were identified using BWA-MEM v0.7.12-r1039 [63, 64]. SAMtools v0.1.19 [65] was used to extract SNPs with MAPQ>/=20 and PoPoolation2 v1.201 [66] was used to extract SNPs with a BQ >/=20, min-count 2, min-coverage 10 and max-coverage 2%. Bedtools v2.15.0 [67] *intersect* was used to identify SNPs within CDS (haemonchus_contortus.PREJEB506.WBPS15.annotations.gff3) and the SNP rate for each isolate was calculated by dividing the number of SNPs in the CDS by the number of nucleotides in the CDS. The ‘SNP rate difference’ was calculated as the MHco18 SNP rate minus the MHco3 SNP rate.

### Gene enrichment analysis

Putative *C. elegans* homologues were identified with BLASTP (*C. elegans* WS264 proteins, e-value < 0.001) and tissue, phenotype and gene ontology enrichment analysis was performed with WormBase Enrichment Analysis [68], which uses a hypergeometric test and Benjamini-Hochberg correction of false discovery rate (Q < 0.1 for significance).

### Data visualisation

Upset plots were generated with UpSetR v1.4.0 [69] and genome-wide expression and *F*_ST_ plots were generated with KaryoploteR v1.8.8 [70]. All other plots were generated with ggplot2 v3.2.1 [71].

### Data availability

Fastq files of trimmed reads have been submitted to ENA under study accession PRJEB4207. The *H. contortus* V4 genome and annotations (version WBPS15) are available at WormBase Parasite [72, 73] https://parasite.wormbase.org/Haemonchus_contortus_prjeb506.

### Real-time quantitative PCR (RT-qPCR)

Total RNA was extracted from 20 male or female worms from each of three different donor sheep per isolate (MHco3 and MHco18). Total RNA (1 μg) was used for oligo(dT) primed cDNA synthesis (SuperScript® III First-Strand Synthesis System, ThermoFisher, 18080051), with no-reverse transcriptase controls included for each sample. cDNA was diluted 1:10 for RT-qPCR (other than for *HCON_00155390:cky-1* in female samples, where neat cDNA was used due to the low abundance of this transcript) and 1 μl added to each reaction. RT-qPCR was undertaken following the Brilliant III Ultra Fast SYBR QPCR Master Mix protocol (Agilent Technologies, 600882) and results were analysed with MxPro v4.10. Gene expression was normalised to *β-actin* (*HCON_00135080*). Primer sequences are listed in Table S7.

## Acknowledgements

This work was funded by a Biotechnology and Biological Sciences Research Council (BBSRC) strategic Lola [BB/M003949], Scottish Government’s Rural and Environment Science and Analytical Services (RESAS) Division and by the Wellcome Trust [206194]. RL is supported by a Wellcome Clinical Research Career Development Fellowship [216614/Z/19/Z], SRD is supported by a UKRI Future Leaders Fellowship [MR/T020733/1]. We thank Professor John Gilleard for access to RNA-seq datasets for MHco4 and MHco10 isolates and Dr Bruce Rosa for advice and sharing scripts for the DESEQ2 batch correction. For the purpose of Open Access, the author has applied a CC BY public copyright licence to any Author Accepted Manuscript version arising from this submission.

## Supplementary Figure Legends

**Figure S1**. Study design. A. The *Haemonchus contortus* MHco3 isolate is fully drug susceptible and the MHco18 isolate is multi-drug resistant; these are the two parental populations used to generate the genetic cross, of which the parent and F2 adult populations are used in this study. One donor sheep was infected per population. Nomenclature, drug treatment, expected resistance phenotype(s) and adult worm samples used in our analyses are described. B. Pairwise comparisons of differential gene expression included in each analysis. For each analysis, genes were filtered for significant differential expressed (adjusted P<0.01) in all pairwise comparisons marked ✓ and for non-significant differential expression (adjusted P>0.01) in all pairwise comparisons marked ✗. Pairwise comparisons marked - were not included in the analysis. Genes with significant differential expression were filtered to retain only those where the direction of log fold change (>0 or <0) was the same in all pairwise comparisons i.e. for inclusion, a gene could not be upregulated in a resistant population in one pairwise comparison and downregulated in a resistant population in another.

**Figure S2**. Sample QC. A. Total reads sequenced for each sample. B. Percentage of reads mapped to MHco3 reference genome for each strain.

**Figure S3**. Heatmap of female samples clustered by the top 500 genes based on row variance in the regularised log transformation. Two clear clusters appear: the F2CTL samples plus sample 1 of every other group (left branch) and all other samples (right branch).

**Figure S4**. Upset plots showing the number of shared upregulated (A) and downregulated (B) genes in different pairwise comparisons for male (M) and female (F) samples.

**Figure S5**. Genome-wide karyoplots showing genomic loci of genes with significant upregulation (yellow) or downregulation (blue). Point size corresponds to significance. A. MHco18 vs MHco3 males, B. MHco18 vs MHco3 females, C. F2IVM vs MHco3 males, D. F2IVM vs MHco3 females and E. F2IVM vs F2CTL males. Panel F shows genetic differentiation (*F*_ST_) between the F3 generation of the genetic cross pre- and post-ivermectin selection [18].

**Figure S6**. Normalised read counts for all male samples for putative candidate ivermectin target and/or resistance genes from the literature.

**Figure S7**. Normalised read counts for all female samples for putative candidate ivermectin target and/or resistance genes from the literature.

**Figure S8**. GO, phenotype and tissue enrichment analysis of *C. elegans* homologues of differentially expressed *H. contortus* genes.

**Figure S9**. Scatter plots showing SNP rate versus differential expression in males (A) and females (B). Points represent genes that are differentially expressed in the parental isolates: light blue if differentially expressed in MHco18 vs MHco3 only, dark blue if also differentially expressed in F2IVM vs F2CTL and F2IVM vs MHco3 (males) or F2IVM vs MHco3 (females).

## References

1. Mectizan Donation Program. https://mectizan.org, (accessed 18.12.2020).

2. Kaplan, R.M. (2004) Drug resistance in nematodes of veterinary importance: a status report. Trends Parasitol 20 (10), 477–81.

3. Kaplan, R.M. and Vidyashankar, A.N. (2012) An inconvenient truth: global worming and anthelmintic resistance. Vet Parasitol 186 (1-2), 70–8.

4. Doyle, S.R. et al. (2017) Genome-wide analysis of ivermectin response by Onchocerca volvulus reveals that genetic drift and soft selective sweeps contribute to loss of drug sensitivity. PLoS Negl Trop Dis 11 (7), e0005816.

5. Osei-Atweneboana, M.Y. et al. (2011) Phenotypic evidence of emerging ivermectin resistance in Onchocerca volvulus. PLoS Negl Trop Dis 5 (3), e998.

6. Osei-Atweneboana, M.Y. et al. (2007) Prevalence and intensity of Onchocerca volvulus infection and efficacy of ivermectin in endemic communities in Ghana: a two-phase epidemiological study. Lancet 369 (9578), 2021–9.

7. Williamson, S.M. et al. (2011) Candidate anthelmintic resistance-associated gene expression and sequence polymorphisms in a triple-resistant field isolate of Haemonchus contortus. Mol Biochem Parasitol 180 (2), 99–105.

8. Dicker, A.J. et al. (2011) Gene expression changes in a P-glycoprotein (Tci-pgp-9) putatively associated with ivermectin resistance in Teladorsagia circumcincta. Int J Parasitol 41 (9), 935–42.

9. El-Abdellati, A. et al. (2011) Altered avr-14B gene transcription patterns in ivermectin-resistant isolates of the cattle parasites, Cooperia oncophora and Ostertagia ostertagi. Int J Parasitol 41 (9), 951–7.

10. Martinez-Valladares, M. et al. (2012) Teladorsagia circumcincta: Molecular characterisation of the avr-14B subunit and its relatively minor role in ivermectin resistance. Int J Parasitol Drugs Drug Resist 2, 154–61.

11. Kotze, A.C. et al. (2014) Recent advances in candidate-gene and whole-genome approaches to the discovery of anthelmintic resistance markers and the description of drug/receptor interactions. Int J Parasitol Drugs Drug Resist 4 (3), 164–84.

12. Blaxter, M. and Koutsovoulos, G. (2015) The evolution of parasitism in Nematoda. Parasitology 142 Suppl 1, S26–39.

13. International Helminth Genomes, C. (2018) Comparative genomics of the major parasitic worms. Nat Genet.

14. Doyle, S.R. et al. (2020) Genomic and transcriptomic variation defines the chromosome-scale assembly of Haemonchus contortus, a model gastrointestinal worm. Commun Biol 3 (1), 656.

15. Doyle, S.R. et al. (2018) A Genome Resequencing-Based Genetic Map Reveals the Recombination Landscape of an Outbred Parasitic Nematode in the Presence of Polyploidy and Polyandry. Genome Biol Evol 10 (2), 396–409.

16. Salle, G. et al. (2019) The global diversity of Haemonchus contortus is shaped by human intervention and climate. Nat Commun 10 (1), 4811.

17. Doyle, S.R. et al. (2019) Population genomic and evolutionary modelling analyses reveal a single major QTL for ivermectin drug resistance in the pathogenic nematode, Haemonchus contortus. BMC Genomics 20 (1), 218.

18. Doyle, S.R. et al. (2021) Genomic landscape of drug response reveals novel mediators of anthelmintic resistance. bioRxiv, 2021.11.12.465712.

19. Packer, J.S. et al. (2019) A lineage-resolved molecular atlas of C. elegans embryogenesis at single-cell resolution. Science 365 (6459).

20. Nollen, E.A. et al. (2004) Genome-wide RNA interference screen identifies previously undescribed regulators of polyglutamine aggregation. Proc Natl Acad Sci U S A 101 (17), 6403–8.

21. Lin, Y. et al. (2008) Activity-dependent regulation of inhibitory synapse development by Npas4. Nature 455 (7217), 1198–204.

22. Ooe, N. et al. (2009) Functional characterization of basic helix-loop-helix-PAS type transcription factor NXF in vivo: putative involvement in an “on demand” neuroprotection system. J Biol Chem 284 (2), 1057–63.

23. Menez, C. et al. (2019) The transcription factor NHR-8: A new target to increase ivermectin efficacy in nematodes. PLoS Pathog 15 (2), e1007598.

24. Khan, S. et al. (2020) A Whole Genome Re-Sequencing Based GWA Analysis Reveals Candidate Genes Associated with Ivermectin Resistance in Haemonchus contortus. Genes (Basel) 11 (4).

25. Le Jambre, L.F. et al. (1995) Characterisation of an avermectin resistant strain of Australian Haemonchus contortus. Int J Parasitol 25 (6), 691–8.

26. Le Jambre, L.F. et al. (2000) Inheritance of avermectin resistance in Haemonchus contortus. Int J Parasitol 30 (1), 105–11.

27. Janssen, I.J. et al. (2015) Transgenically expressed Parascaris P-glycoprotein-11 can modulate ivermectin susceptibility in Caenorhabditis elegans. Int J Parasitol Drugs Drug Resist 5 (2), 44–7.

28. Yan, R. et al. (2012) The role of several ABC transporter genes in ivermectin resistance in Caenorhabditis elegans. Vet Parasitol 190 (3-4), 519–29.

29. Jospin, M. et al. (2007) UNC-80 and the NCA ion channels contribute to endocytosis defects in synaptojanin mutants. Curr Biol 17 (18), 1595–600.

30. Yeh, E. et al. (2008) A putative cation channel, NCA-1, and a novel protein, UNC-80, transmit neuronal activity in C. elegans. PLoS Biol 6 (3), e55.

31. Altun, Z.F., Herndon, L.A., Wolkow, C.A., Crocker, C., Lints, R. and Hall D.H. WormAtlas. http://www.wormatlas.org, (accessed 24.03.21).

32. Nath, R.D. et al. (2016) C. elegans Stress-Induced Sleep Emerges from the Collective Action of Multiple Neuropeptides. Curr Biol 26 (18), 2446–2455.

33. Hill, A.J. et al. (2014) Cellular stress induces a protective sleep-like state in C. elegans. Curr Biol 24 (20), 2399–405.

34. Rezansoff, A.M. et al. (2019) The confounding effects of high genetic diversity on the determination and interpretation of differential gene expression analysis in the parasitic nematode Haemonchus contortus. Int J Parasitol 49 (11), 847–858.

35. Han, B. et al. (2015) An evolutionarily conserved switch in response to GABA affects development and behavior of the locomotor circuit of Caenorhabditis elegans. Genetics 199 (4), 1159–72.

36. Steel, J.W. (1993) Pharmacokinetics and metabolism of avermectins in livestock. Vet Parasitol 48 (1-4), 45–57.

37. Hisamoto, N. et al. (2016) The C. elegans Discoidin Domain Receptor DDR-2 Modulates the Met-like RTK-JNK Signaling Pathway in Axon Regeneration. PLoS Genet 12 (12), e1006475.

38. Matsuo, K. et al. (2019) Visualization of endogenous NID-1 and EMB-9 in C. elegans. MicroPubl Biol 2019.

39. Branicky, R. et al. (2014) The voltage-gated anion channels encoded by clh-3 regulate egg laying in C. elegans by modulating motor neuron excitability. J Neurosci 34 (3), 764–75.

40. Doyle, S.R. and Cotton, J.A. (2019) Genome-wide Approaches to Investigate Anthelmintic Resistance. Trends Parasitol 35 (4), 289–301.

41. Kim, D. et al. (2015) HISAT: a fast spliced aligner with low memory requirements. Nat Methods 12 (4), 357–60.

42. Kim, D. et al. (2019) Graph-based genome alignment and genotyping with HISAT2 and HISAT-genotype. Nat Biotechnol 37 (8), 907–915.

43. Cheeseman, C.L. et al. (2001) High-affinity ivermectin binding to recombinant subunits of the Haemonchus contortus glutamate-gated chloride channel. Mol Biochem Parasitol 114 (2), 161–8.

44. Dent, J.A. et al. (2000) The genetics of ivermectin resistance in Caenorhabditis elegans. Proc Natl Acad Sci U S A 97 (6), 2674–9.

45. Dent, J.A. et al. (1997) avr-15 encodes a chloride channel subunit that mediates inhibitory glutamatergic neurotransmission and ivermectin sensitivity in Caenorhabditis elegans. EMBO J 16 (19), 5867–79.

46. Laing, S.T. et al. (2012) The transcriptional response of Caenorhabditis elegans to Ivermectin exposure identifies novel genes involved in the response to reduced food intake. PLoS One 7 (2), e31367.

47. Scott, E.W. et al. (1991) Fecundity of anthelmintic resistant adult Haemonchus contortus after exposure to ivermectin or benzimidazoles in vivo. Res Vet Sci 50 (2), 247–9.

48. Laing, R. et al. (2017) Ivermectin -Old Drug, New Tricks? Trends Parasitol 33 (6), 463–472.

49. Redman, E. et al. (2008) Genetics of mating and sex determination in the parasitic nematode Haemonchus contortus. Genetics 180 (4), 1877–87.

50. Matouskova, P. et al. (2018) UDP-glycosyltransferase family in Haemonchus contortus: Phylogenetic analysis, constitutive expression, sex-differences and resistance-related differences. Int J Parasitol Drugs Drug Resist 8 (3), 420–429.

51. Vokral, I. et al. (2012) The metabolism of flubendazole and the activities of selected biotransformation enzymes in Haemonchus contortus strains susceptible and resistant to anthelmintics. Parasitology 139 (10), 1309–16.

52. Laing, S.T. et al. (2010) Characterization of the xenobiotic response of Caenorhabditis elegans to the anthelmintic drug albendazole and the identification of novel drug glucoside metabolites. Biochem J 432 (3), 505–14.

53. Stasiuk, S.J. et al. (2019) Similarities and differences in the biotransformation and transcriptomic responses of Caenorhabditis elegans and Haemonchus contortus to five different benzimidazole drugs. Int J Parasitol Drugs Drug Resist 11, 13–29.

54. Roos, M.H. et al. (2004) Genetic analysis of inbreeding of two strains of the parasitic nematode Haemonchus contortus. Int J Parasitol 34 (1), 109–15.

55. Martin, M. (2011) Cutadapt removes adapter sequences from high-throughput sequencing reads. EMBnet.journal 17 (1), 10–12.

56. Joshi, N.F. JN, Sickle: A sliding-window, adaptive, quality-based trimming tool for FastQ files (Version 1.33), 2011.

57. Afgan, E. et al. (2018) The Galaxy platform for accessible, reproducible and collaborative biomedical analyses: 2018 update. Nucleic Acids Res 46 (W1), W537–W544.

58. RStudioTeam, RStudio: Integrated Development for R, 2016.

59. Andrews, S., FastQC A Quality Control tool for High Throughput Sequence Data.

60. Ewels, P. et al. (2016) MultiQC: summarize analysis results for multiple tools and samples in a single report. Bioinformatics 32 (19), 3047–8.

61. Liao, Y. et al. (2014) featureCounts: an efficient general purpose program for assigning sequence reads to genomic features. Bioinformatics 30 (7), 923–30.

62. Love, M.I. et al. (2014) Moderated estimation of fold change and dispersion for RNA-seq data with DESeq2. Genome Biol 15 (12), 550.

63. Li, H. and Durbin, R. (2009) Fast and accurate short read alignment with Burrows-Wheeler transform. Bioinformatics 25 (14), 1754–60.

64. Li, H. (2013) Aligning sequence reads, clone sequences and assembly contigs with BWA-MEM. arXiv.org.

65. Li, H. et al. (2009) The Sequence Alignment/Map format and SAMtools. Bioinformatics 25 (16), 2078–9.

66. Kofler, R. et al. (2011) PoPoolation: a toolbox for population genetic analysis of next generation sequencing data from pooled individuals. PLoS One 6 (1), e15925.

67. Quinlan, A.R. and Hall, I.M. (2010) BEDTools: a flexible suite of utilities for comparing genomic features. Bioinformatics 26 (6), 841–2.

68. Angeles-Albores, D. et al. (2016) Tissue enrichment analysis for C. elegans genomics. BMC Bioinformatics 17 (1), 366.

69. Conway, J.R. et al. (2017) UpSetR: an R package for the visualization of intersecting sets and their properties. Bioinformatics 33 (18), 2938–2940.

70. Gel, B. and Serra, E. (2017) karyoploteR: an R/Bioconductor package to plot customizable genomes displaying arbitrary data. Bioinformatics 33 (19), 3088–3090.

71. Wickham, H. (2016) ggplot2: Elegant Graphics for Data Analysis, Springer-Verlag New York.

72. Bolt, B.J. et al. (2018) Using WormBase ParaSite: An Integrated Platform for Exploring Helminth Genomic Data. Methods Mol Biol 1757, 471–491.

73. Howe, K.L. et al. (2017) WormBase ParaSite - a comprehensive resource for helminth genomics. Mol Biochem Parasitol 215, 2–10.

