## Supplementary Figures for "Transcriptomic analyses implicate neuronal plasticity and chloride homeostasis in ivermectin resistance and recovery in a parasitic nematode"

A

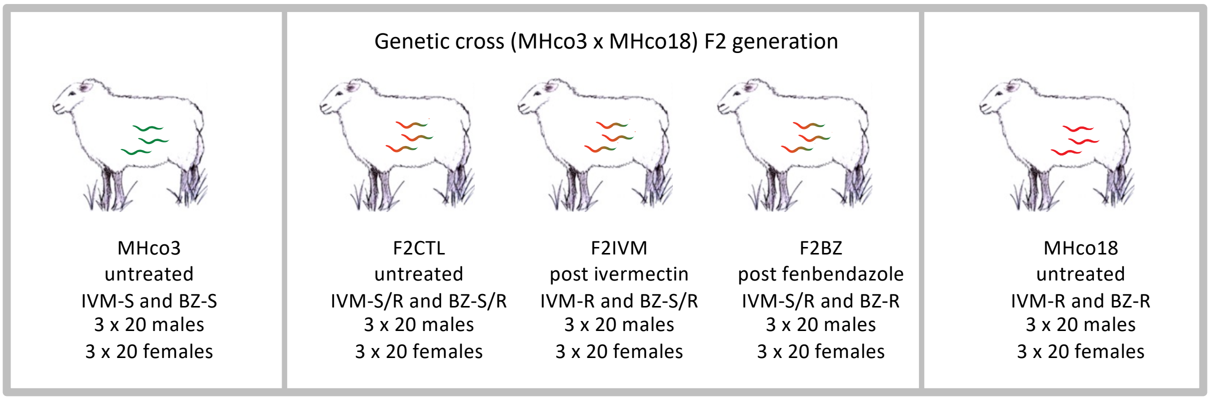


B

| Analysis \ Pairwise comparison | MHco18 v MHco3 males | MHco18 v MHco3 females | F2IVM v MHco3 males | F2IVM v MHco3 females | F2IVM v F2CTL males | F2BZ v MHco3 males | F2BZ v MHco3 females | F2BZ v F2CTL males | MHco4 v MHco3 females | MHco10 v MHco3 females |
|---|---|---|---|---|---|---|---|---|---|---|
| Transcriptomic differences associated with ivermectin resistance | ✓ | ✓ | ✓ | ✓ | ✓ | - | - | - | - | - |
| Differences in gene expression in ivermectin resistant male worms | ✓ | ✕ | ✓ | ✕ | ✓ | - | - | - | - | - |
| Differences in gene expression in ivermectin resistant female worms | ✕ | ✓ | ✕ | ✓ | ✕ | - | - | - | ✓ | ✓ |
| Transcriptomic response to ivermectin exposure | ✕ | ✕ | ✓ | ✓ | ✓ | - | - | - | - | - |
| Similarities in gene expression with benzimidazole selected populations | ✓ | ✓ | ✓ | ✓ | ✓ | ✓ | ✓ | ✓ | - | - |
| Transcriptomic differences associated with benzimidazole resistance | ✓ | ✓ | - | - | - | ✓ | ✓ | ✓ | - | - |

Figure S1. Study design. A. The *Haemonchus contortus* MHco3 isolate is fully drug susceptible and the MHco18 isolate is multi-drug resistant; these are the two parental populations used to generate the genetic cross, of which the parent and F2 adult populations are used in this study. One donor sheep was infected per population. Nomenclature, drug treatment, expected resistance phenotype(s) and adult worm samples used in our analyses are described. B. Pairwise comparisons of differential gene expression included in each analysis. For each analysis, genes were filtered for significant differential expressed (adjusted P<0.01) in all pairwise comparisons marked ✓ and for non-significant differential expression (adjusted P>0.01) in all pairwise comparisons marked ✕. Pairwise comparisons marked - were not included in the analysis. Genes with significant differential expression were filtered to retain only those where the direction of log fold change (>0 or <0) was the same in all pairwise comparisons i.e. for inclusion, a gene could not be upregulated in a resistant population in one pairwise comparison and downregulated in a resistant population in another.

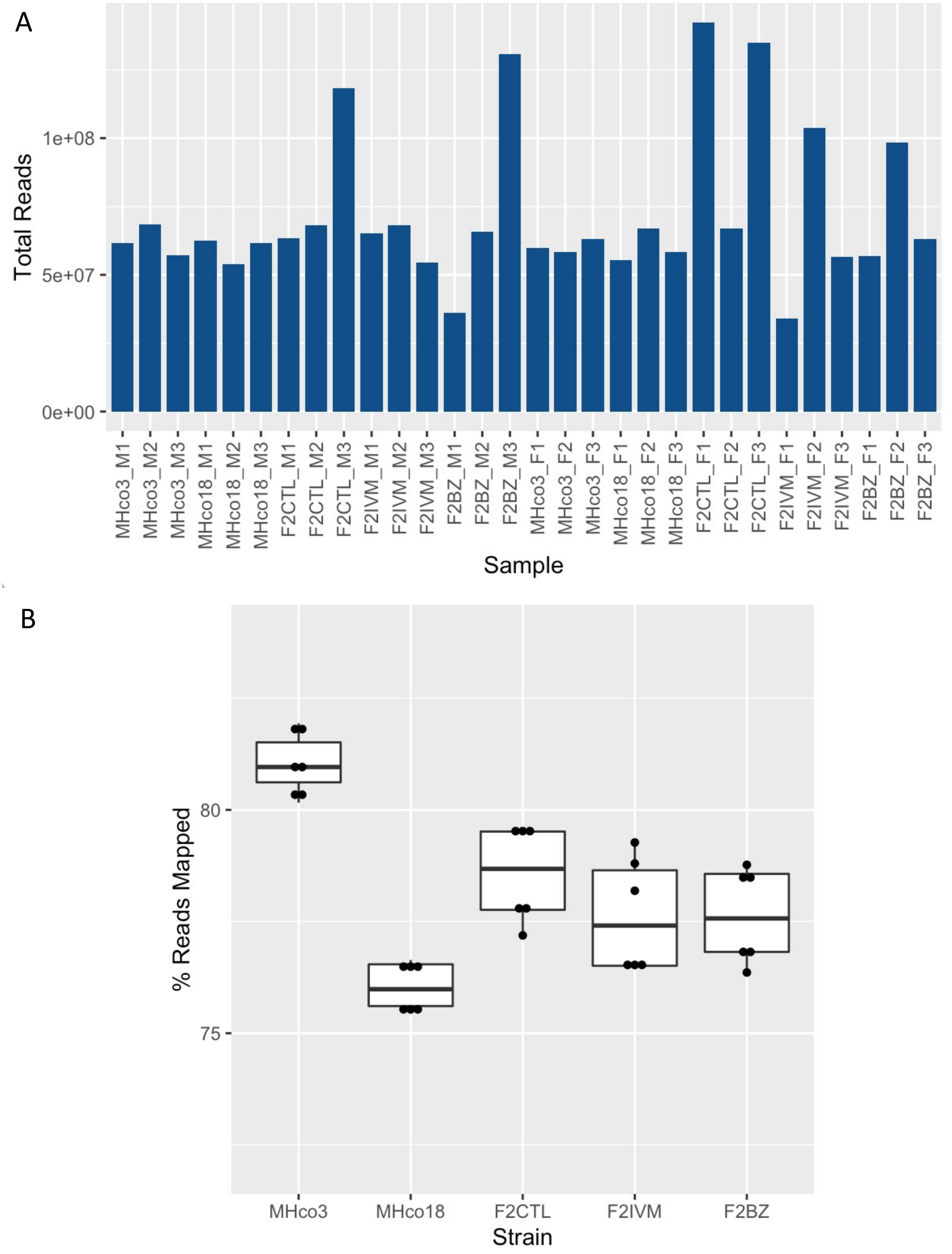


Figure S2. Sample QC. A. Total reads sequenced for each sample. B. Percentage of reads mapped to MHco3 reference genome for each strain.

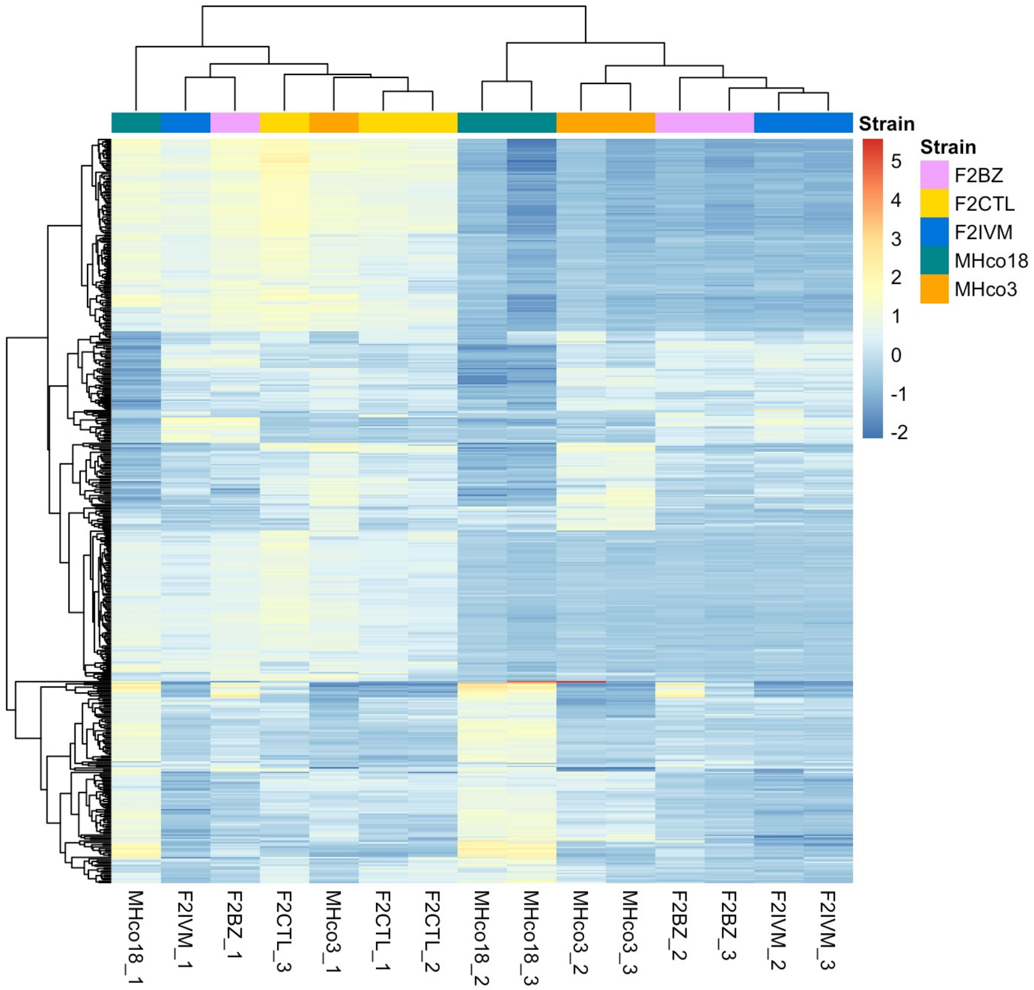


Figure S3. Heatmap of female samples clustered by the top 500 genes based on row variance in the regularised log transformation. Two clear clusters appear: the F2CTL samples plus sample 1 of every other group (left branch) and all other samples (right branch).

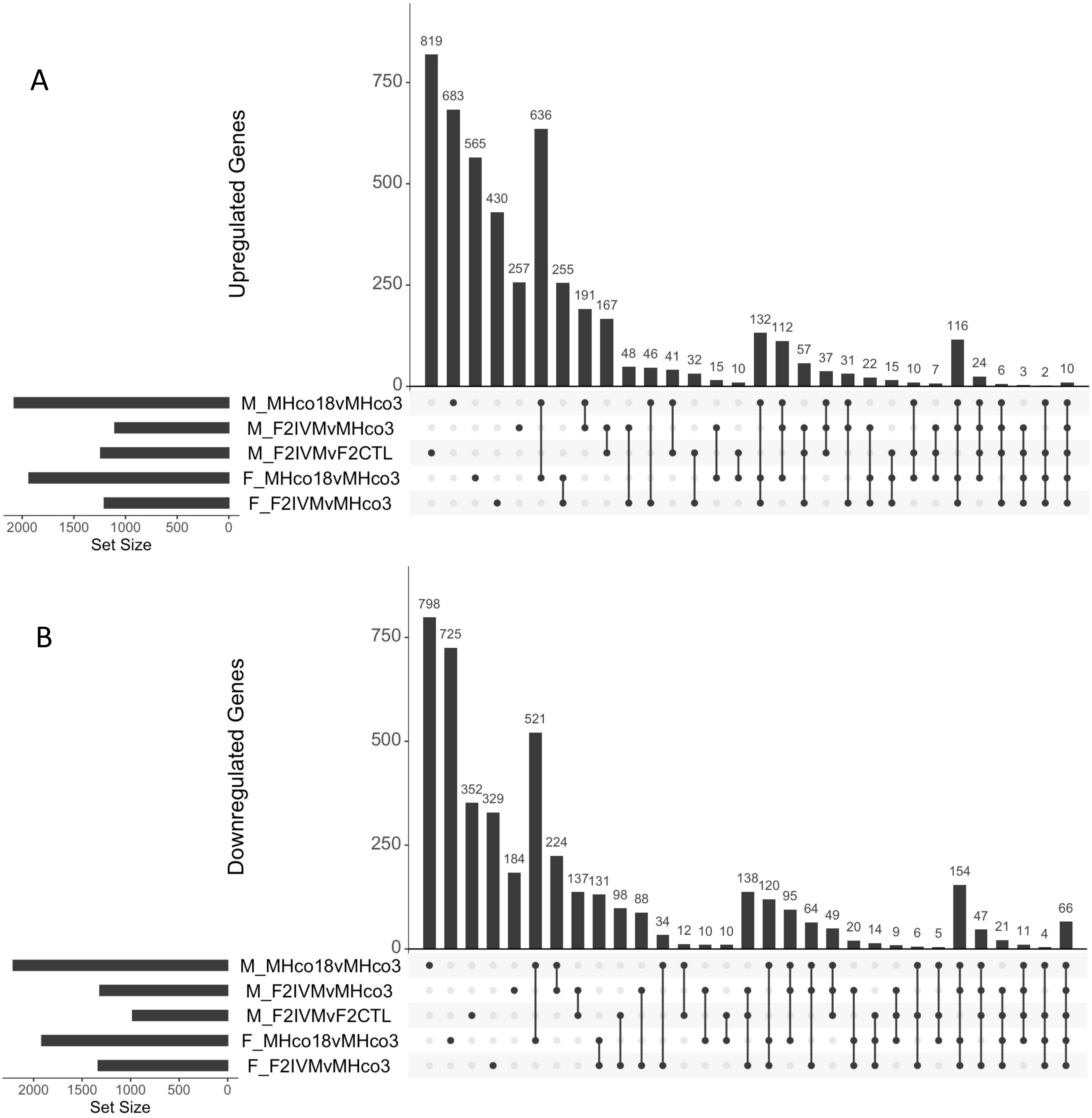


Figure S4. Upset plots showing the number of shared upregulated (A) and downregulated (B) genes in different pairwise comparisons for male (M) and female (F) samples.

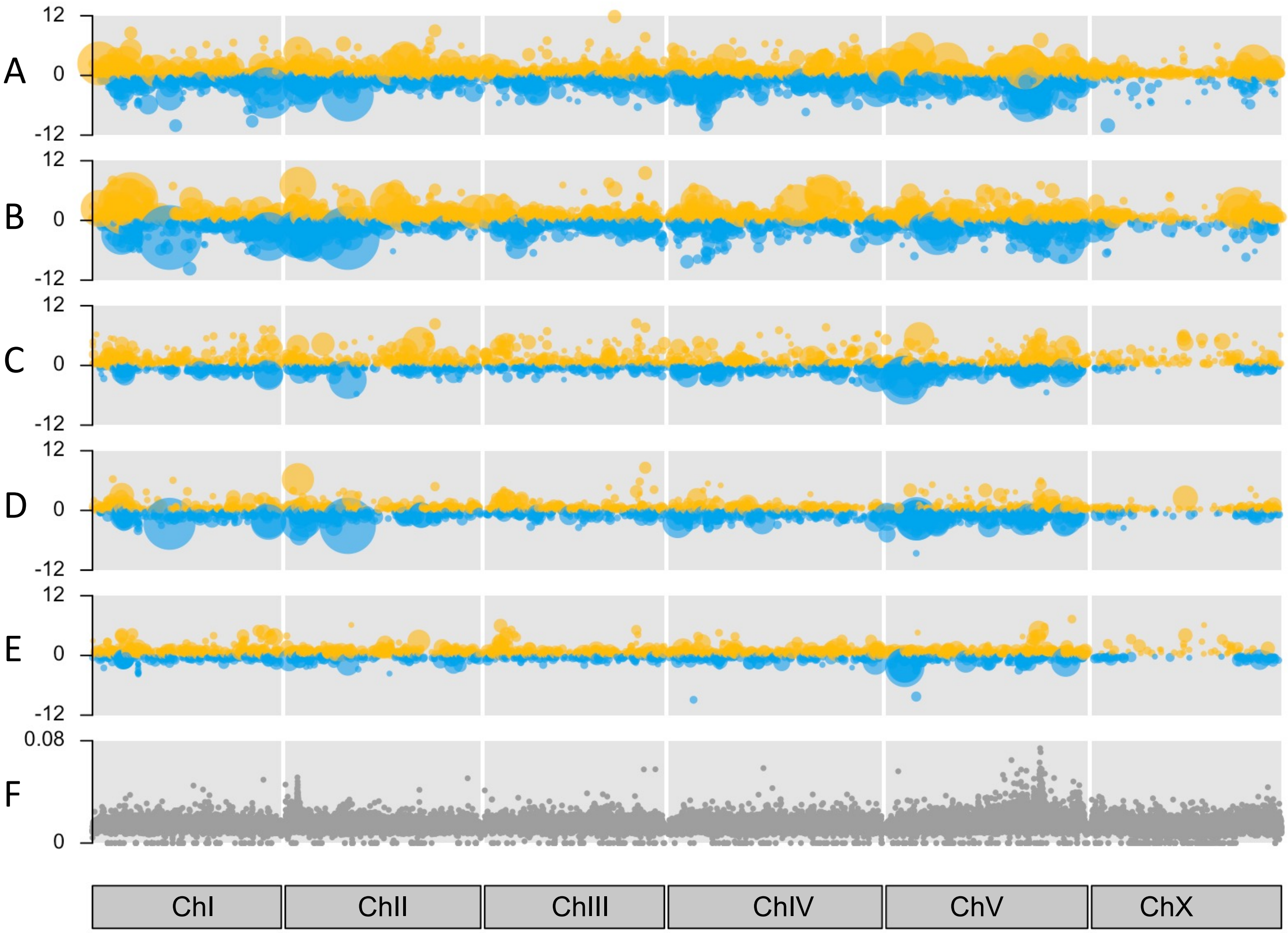


Figure S5. Genome-wide karyoplots showing genomic loci of genes with significant upregulation (yellow) or downregulation (blue). Point size corresponds to significance. A. MHco18 vs MHco3 males, B. MHco18 vs MHco3 females, C. F2IVM vs MHco3 males, D. F2IVM vs MHco3 females and E. F2IVM vs F2CTL males. Panel F shows genetic differentiation ($F_{ST}$) between the F3 generation of the genetic cross pre- and post- ivermectin selection [15].

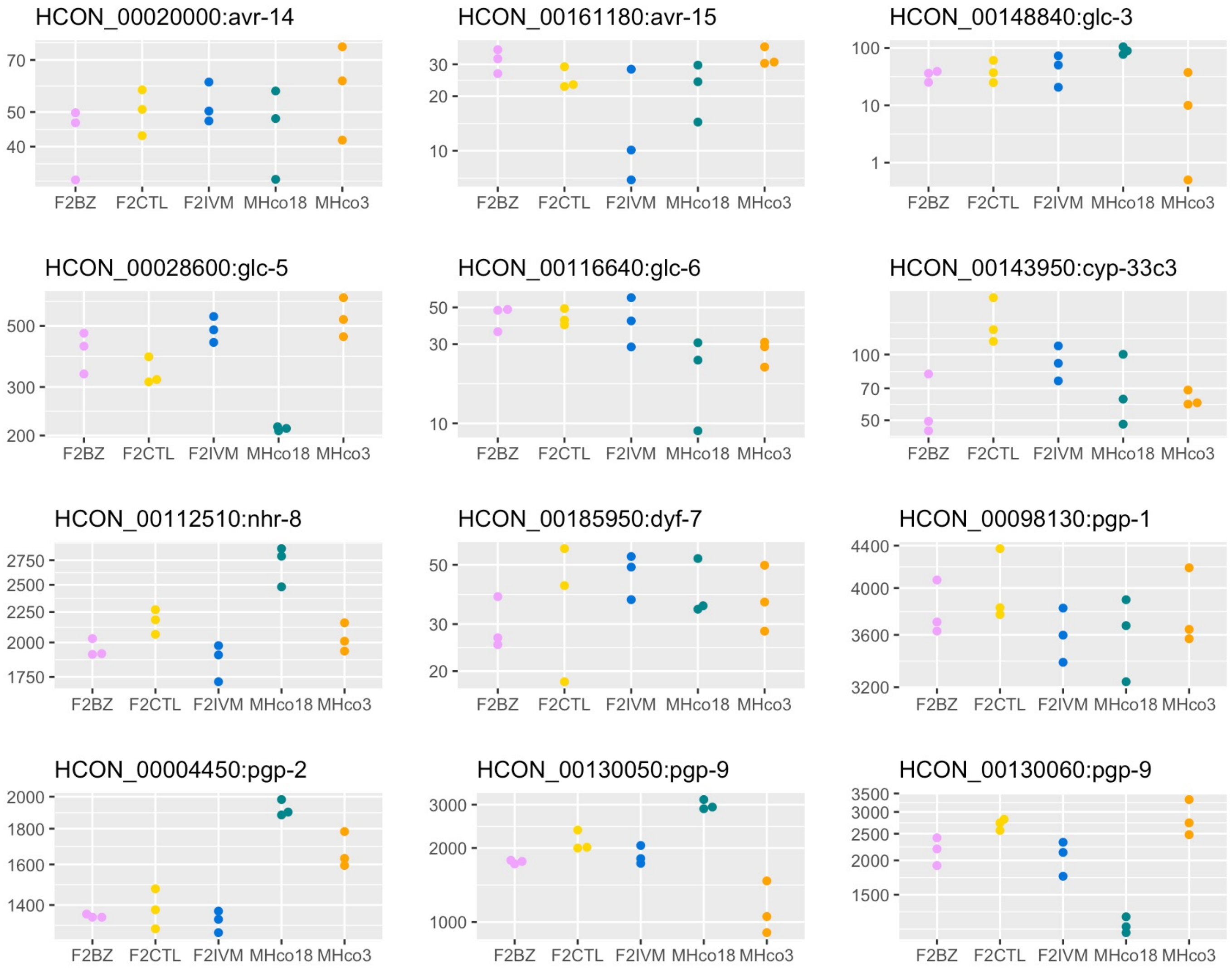


Figure S6. Normalised read counts for all male samples for putative candidate ivermectin target and/or resistance genes from the literature.

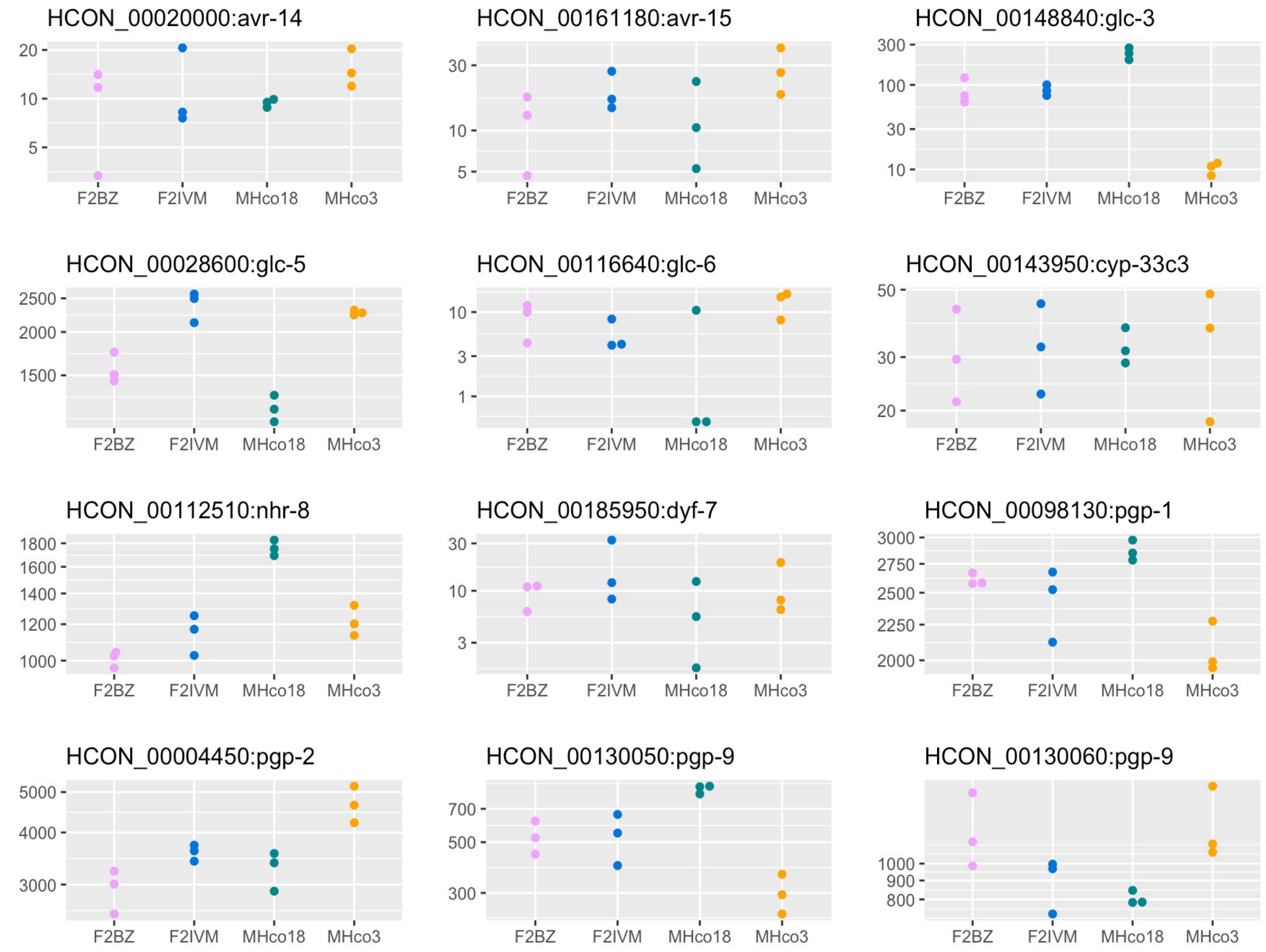


Figure S7. Normalised read counts for all female samples for putative candidate ivermectin target and/or resistance genes from the literature.

Male only

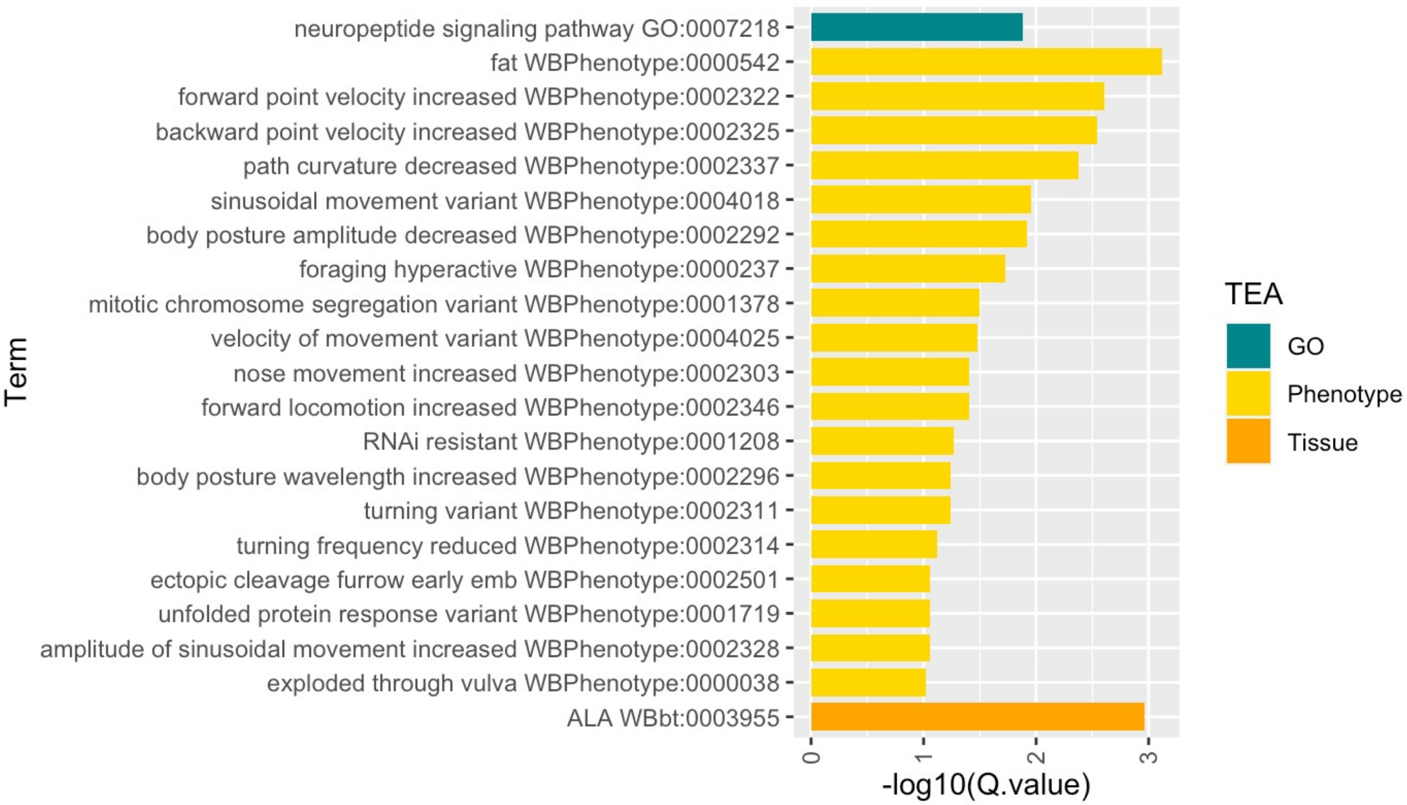


Response to ivermectin treatment

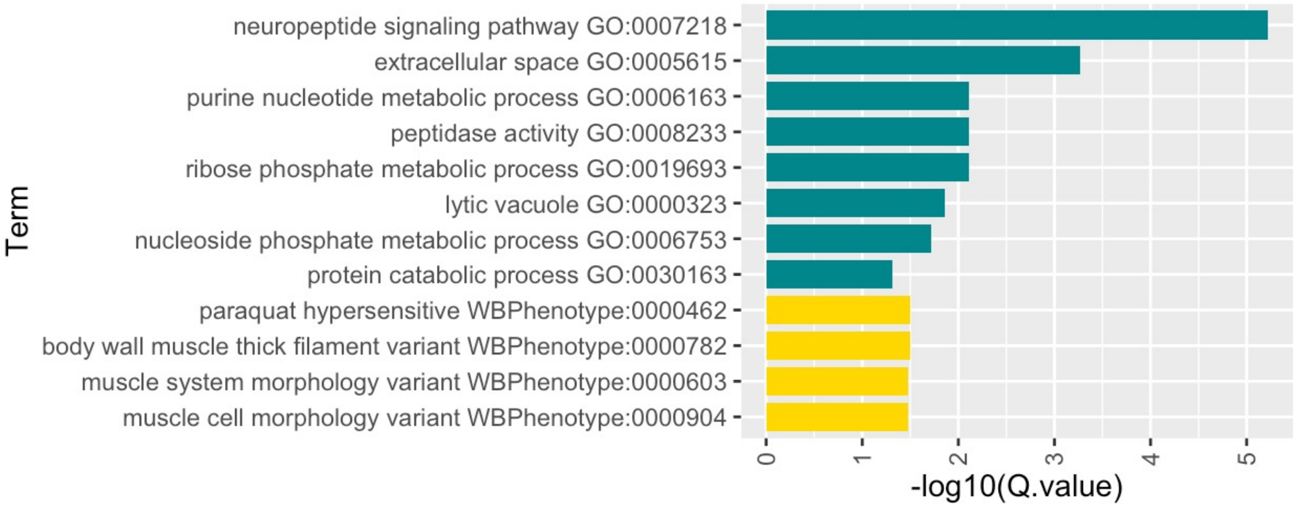


Figure S8. GO, phenotype and tissue enrichment analysis of *C. elegans* homologues of differentially expressed *H. contortus* genes.

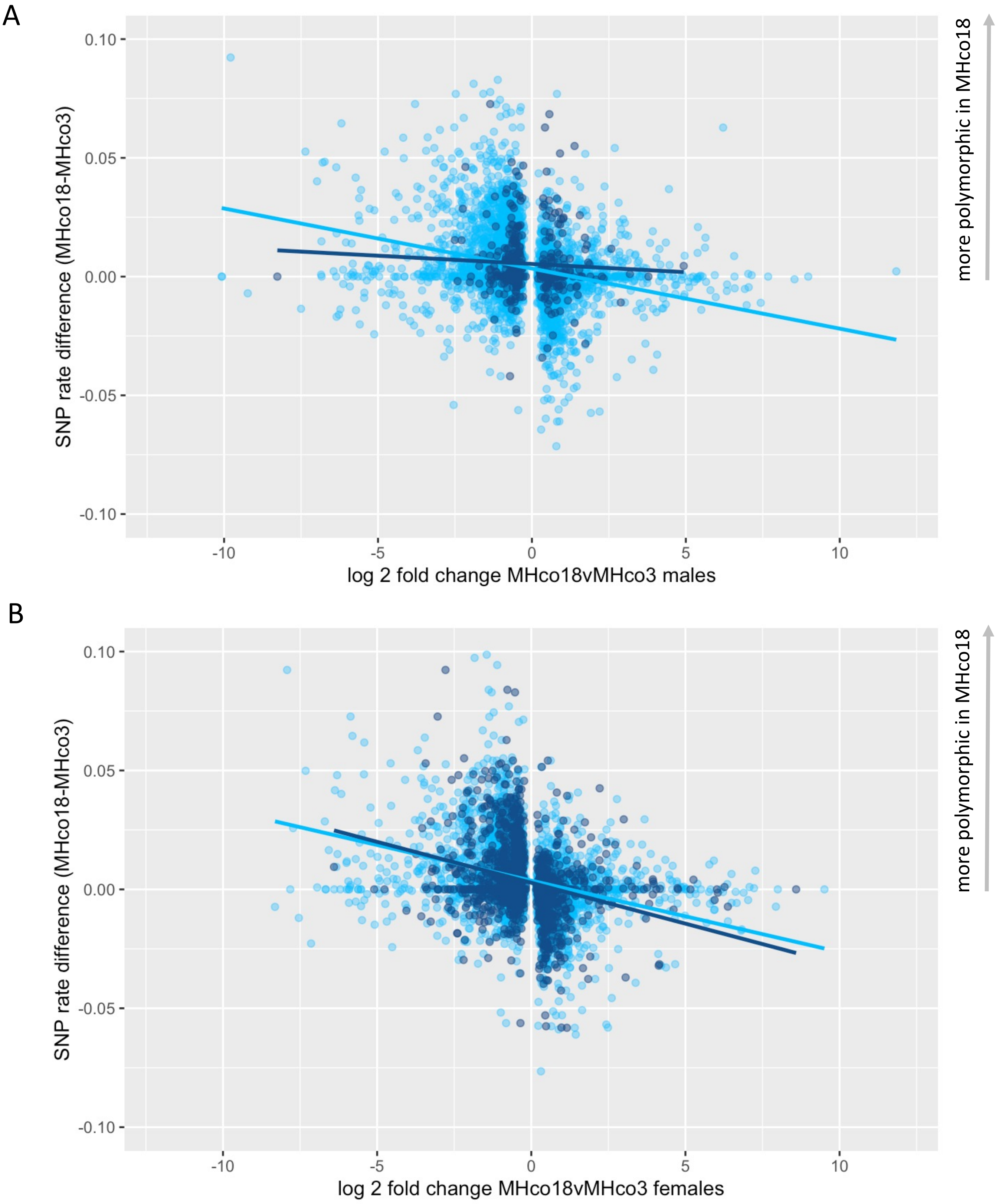


Figure S9. Scatter plots showing SNP rate versus differential expression in males (A) and females (B). Points represent genes that are differentially expressed in the parental isolates: light blue if differentially expressed in MHco18 vs MHco3 only, dark blue if also differentially expressed in F2IVM vs F2CTL and F2IVM vs MHco3 (males) or F2IVM vs MHco3 (females).
